# Sex biased human thymic architecture guides T cell development through spatially defined niches

**DOI:** 10.1101/2023.04.13.536804

**Authors:** Laura N Stankiewicz, Kevin Salim, Emily A Flaschner, Yu Xin Wang, John M Edgar, Bruce ZB Lin, Grace C Bingham, Matthew C Major, Ross D Jones, Helen M Blau, Elizabeth J Rideout, Megan K Levings, Peter W Zandstra, Fabio MV Rossi

**Author notes:** Correspondence (P.W.Z), (F.MV.R). These authors contributed equally (Peter W Zandstra, Fabio MV Rossi).

## Abstract

Within the thymus, regulation of the cellular cross-talk directing T cell development is dependent on spatial interactions within specialized niches. To create a holistic, spatially defined map of tissue niches guiding postnatal T cell development we employed the multidimensional imaging platform CO-detection by indEXing (CODEX), as well as CITE-seq and ATAC-seq. We generated age-matched 4–5-month-old postnatal thymus datasets for male and female donors, and identify significant sex differences in both T cell and thymus biology. We demonstrate a crucial role for JAG ligands in directing thymic-like dendritic cell development, reveal important functions of a novel population of ECM^-^ fibroblasts, and characterize the medullary niches surrounding Hassall’s corpuscles. Together, these data represent a unique age-matched spatial multiomic resource to investigate how sex-based differences in thymus regulation and T cell development arise, and provide an essential resource to understand the mechanisms underlying immune function and dysfunction in males and females.

## Introduction

The thymus is the primary organ responsible for the generation and selection of mature, functional, and self-tolerant T cells^1^. Effective T cell development is a critical component of our immune system and our ability to accurately, and exclusively, identify and kill foreign entities such as pathogens. During postnatal T cell development – the period in life when T cell development is most active^2^ – thymic seeding progenitors migrate to the thymus where they first mature into thymocytes. Thymic architecture is highly organized to provide spatially-defined, stage-specific signaling cues to migrating thymocytes which guide development towards functional mature T cells^3–6^.

Recent single cell sequencing resources demonstrating the diversity of human thymus tissue are incongruous with our current framework of thymus structure and organization^7–19^ which describes a general migratory path a thymocyte takes through the cortex and medulla during conventional αβT cell development. Recent spatial transcriptomic sequencing of human fetal and postnatal thymus has demonstrated that a deeper granularity of thymic niches is present within the tissue and that these niches evolve during fetal development to support different waves of non-conventional T cells^19,20^. However, our understanding of how the granularity and composition of human postnatal thymus niches support conventional and non-conventional T cell development, T-lineage branching decisions, and the development of alternative lineages known to develop in the thymus remains limited^3,4,6^. T cells generated at this stage of postnatal human development will become the foundation of our immune system, patrolling the body for decades^21^. Thus, insights into early postnatal thymus niche biology are crucial to our understanding of how this part of our adaptive immune system is built and how perturbations in postnatal T cell development may emerge as immune dysfunction later in life.

To create a holistic, spatially defined map of tissue niches guiding human postnatal T cell development, alternative lineage development, and age-related changes to thymic architecture, we employed multi dimensional spatial proteomic imaging using CO-detection by inDEXing (CODEX)^22,23^, single cell transcriptomic-proteomic profiling using Cellular Indexing of Transcriptomes and Epitopes (CITE-seq)^24^, and single cell Assay for Transposase Accessible Chromatin (ATAC-seq)^25^. Given the emerging recognition of sex differences in thymus gene expression and function^26–31^, we collected and analysed samples from both male and female donors. Our detailed analysis reveals significant sex differences during early postnatal development that affect both T cell and thymus biology through common and cell type specific mechanisms. Common differences in thymic cells include higher levels of expression of mRNAs associated with energy regulation, translation, and antigen-presentation pathways in female-derived stromal, epithelial, and T cells. Male thymic cells have significantly higher levels of genes associated with adipogenesis, proinflammatory signaling, and ECM pathways. Cell type-specific differences include meaningful thymus-specific sex differences in key cell types and developmental niches which could significantly affect fundamental processes in the development and training of T cells. Finally, we highlight key cell types contributing to thymic involution that exhibit important sex-based differences in thymic growth and early transition towards adipogenesis. These data suggest that differences in the kinetics of thymic involution are present between sexes and, importantly, that mechanisms driving thymic involution begin early in life. Altogether, these data represent a powerful age-matched spatial multiomic resource to investigate how sex-based differences in thymus biology and T cell development arise, and how they contribute to sex differences in diseases caused by immune dysfunction later in life.

## Results

### Spatial multiomic profiling of human postnatal thymus identifies sex-based differences in T cells and thymus biology

We performed CITE-seq, ATAC-seq, and CODEX imaging on human postnatal thymuses ranging from 4 to 33 months of age, including six (3 female, 3 male) 4-5 month old age-matched early postnatal samples (Sup. Table 1), representing the stage of human development where naïve T cell generation is at its highest. Upon receipt, samples were split and processed simultaneously for CODEX imaging or sequencing (Figure 1A). Prior to sequencing we specifically enriched for CD45^-^ non-hematopoietic cells and CD25^+^CD8^-^ regulatory T (Treg) cells to ensure coverage of low abundance cell types found within the thymus. After quality control, we obtained a total of 74,334 cells with CITE-seq, including 19,434 non-T lineage cells, and captured 25,717 nuclei with ATAC-seq. During sequencing sample preparation, we included a comprehensive human universal antibody panel consisting of 137 CITE-seq antibodies (Sup. Table 2), allowing us to compare specific differences in epigenomic, transcriptomic, and proteomic expression kinetics across developing thymocytes, as well as enable direct comparison of cells identified via phenotypic expression in deep imaging to cells captured via CITE-seq.

**Figure 1.**
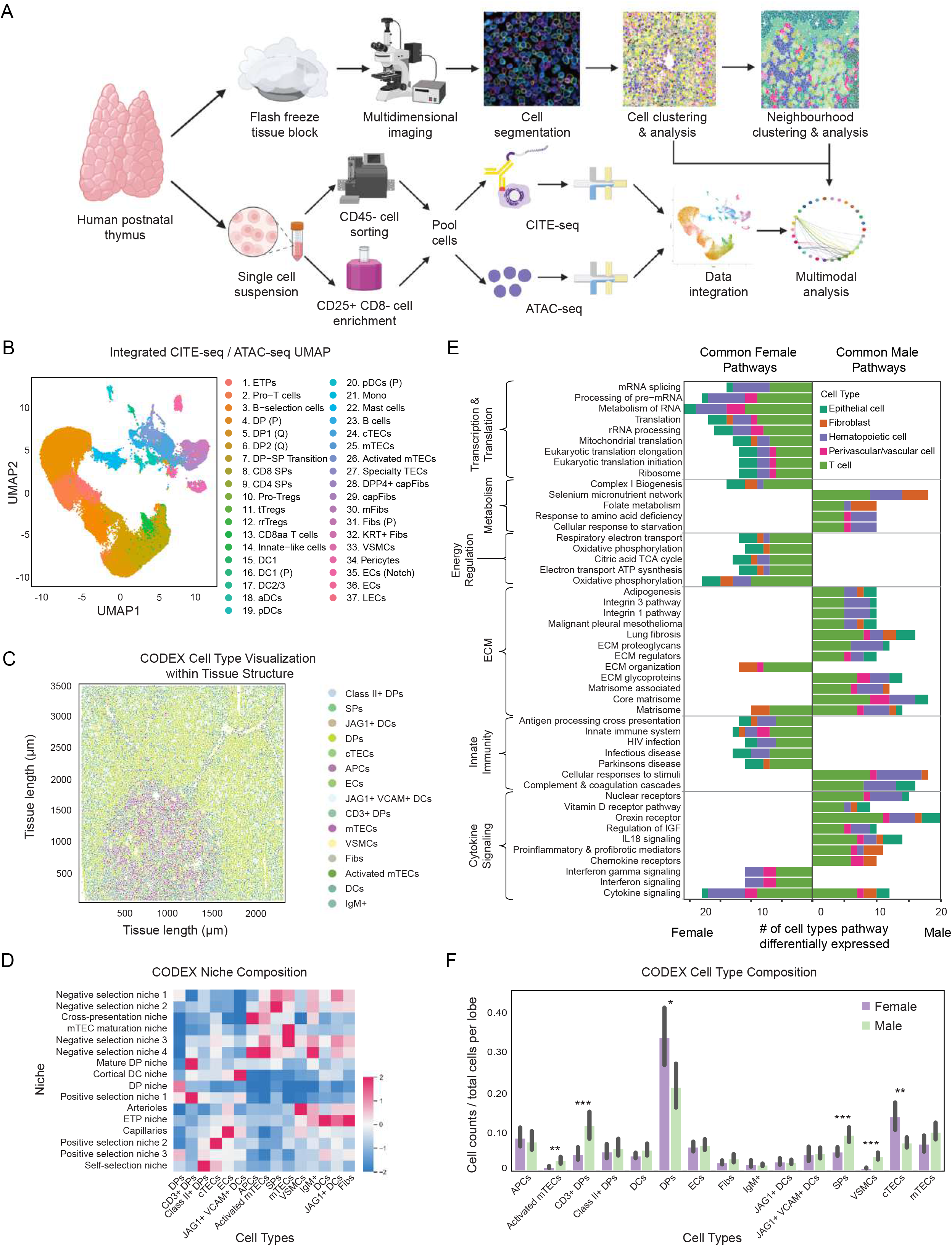
A spatial multi-omic analytical platform identifies sex biased characteristics of thymic niches: A) Graphic describing the experimental design and spatial multiomic analysis. Briefly, human postnatal samples were split and processed simultaneously to capture deep imaging data via CODEX or sequencing data via CITE-seq and ATAC-seq. CODEX data were clustered into proximity-based niches which guided niche analysis of integrated multimodal sequencing data. B) Integrated CITE-seq/ATAC-seq UMAP displaying the 37 cell types captured via sequencing. C) Representative visualization of cell clusters captured via CODEX within the thymic lobule tissue structure. Cell clusters were named based on marker gene expression and spatial localization within the tissue. D) Heat map depicting proximity based niche composition probabilities captured via CODEX. Positive values depict cell types which are frequently found in that niche. E) Histogram showing the combined top 25 common GSEA pathways upregulated in male thymic cells and top 25 pathways commonly upregulated in female thymic cells. Pathways represent common sex-based differences among male and female cell types and are broken down by cell type category to show distribution across cell types (p_adjust_<0.05). F) Bar plot showing differences in normalized cell counts per cell type identified via CODEX. Cell type counts are normalized to the total number of cells in their respective imaged lobule; * p<0.05, ** p<0.01, *** p<0.001.

Cells captured with CITE-seq were clustered based on transcriptional expression, annotated based on marker gene and surface protein expression (Sup. Figure 1A), and aligned where possible with existing human thymus datasets (Sup. Table 3)^7,8^. ATAC-seq clusters were computationally labeled using our 37 CITE-seq clusters as reference, which identified 34 ATAC-seq cluster labels to be transferred prior to dataset integration (Figure 1B, Sup. Figure 1B). We captured 54,900 thymocytes spanning development from early thymic progenitors (ETPs) to mature single positive (SP) T cells, innate-like cells, CD8aa T cells, and Tregs. For Tregs, we identified three populations which expressed canonical Treg lineage markers, namely Treg progenitors (Pro-Tregs), thymic Tregs (tTregs), and recirculating/resident Tregs (rrTregs)^32^. We also identified various antigen presenting cells, such as B cells, mast cells, monocytes, and six populations of dendritic cells (DCs)^33^. In addition to the activated dendritic cells (aDCs), plasmacytoid dendritic (pDCs), DC1, and DC2/3 populations described by Park et al.^7^, we found proliferating populations of pDCs and DC1. We also captured 7093 epithelial cells, including cortical epithelial cells (cTECs), medullary epithelial cells (mTECs), activated mTECs, and specialty TECs.

Importantly, we enriched and captured 7721 mesenchymal cells, a population that has recently been highlighted for newly described roles in negative selection and thymic involution^9,19, 34–36^. Subclustering of this population revealed several important mesenchymal cell types including two populations of endothelial cells (ECs) defined by differential expression of Notch ligands JAG1, JAG2, DLL1, and DLL4 (ECs, ECs (Notch)). Additionally, we identified lymphatic endothelial cells (LECs), pericytes, vascular smooth muscle cells (VSMCs), and five distinct fibroblast cell types, including DPP4^+^ capsular fibroblasts (DPP4+ capFibs), capsule fibroblasts (capFibs), medullary fibroblasts (mFibs), KRT^+^ fibroblasts (KRT+ Fibs), and unique population of proliferating fibroblasts (Fibs (P)).

We designed a custom 48 antibody panel for CODEX imaging of human postnatal thymus to study the architecture and function of niches guiding thymocyte development, with a specific focus on defining the niche characteristics guiding T-lineage branch points. Stage specific thymocyte phenotyping markers (CD62L, CCR7, CD1A, CD5, CD7, CD4, CD8, CD3, CD45RO, CD45RA, FOXP3, SATB1) identified the location of subsets of interest, such as CD3^+^ double positive cells (DPs) undergoing T-lineage commitment towards CD4 or CD8 T cells. Phenotyping markers for non-T lineage hematopoietic cells (CD19, CD11c, CD11b, CD68), epithelial cells (EPCAM, KRT5/8), mural cells (MCAM, SMA), endothelial cells (CD31), and fibroblasts (PDGFRA) identified the remaining known cell types defining thymic architecture and niche composition. Finally, we included functional markers to define patterns of antigen presentation (CD86), MHC class I and II expression (HLA-ABC, HLA-DR,DP,DQ), adhesion ligands (ICAM, VCAM), Notch ligand expression (DLL1, DLL4, JAG1, JAG2), T cell activation (PD-1, PD-L1), proliferation (Ki67), and enzymatic regulation (15-PDGH). In sum, our deep imaging panel enabled a holistic approach to defining thymic biology and allowed us to investigate the spatially-regulated mechanisms directing human T cell development.

Using neural-network driven cell segmentation and Leiden-based clustering^23^ we identified individual cells within thymic tissue for each individual sample (Sup. Figure 1C). We annotated cell types based on location within the tissue and corresponding phenotypic expression compared to our CITE-seq clusters (Figure 1C) and performed proximity-based neighbourhood clustering to identify specific niches as detailed previously^23^. These tissue niches were annotated based on location and cell type composition (Figure 1D; Sup. Figure 1C). This computational analysis allowed us to quantify proximity-based cell-cell interactions (Sup. Figure 1E) and acted as a platform to interrogate spatially-defined thymic niche biology via integrated sequencing-imaging analysis.

We compared our age-matched datasets to identify common and cell type-specific differences in T cell development and thymus biology within early postnatal donors. Because of known sex differences in thymus and T cell gene expression and function^31^, we analysed male and female samples separately. Of the differentially expressed genes only 6% were found on sex chromosomes (Sup. Table 4), in line with prior reports of sex-biased gene expression on autosomes^37–40^. Gene set enrichment analysis (GSEA) on male and female cells for each cell type identified via CITE-seq revealed pathways commonly upregulated in either male or female cells (Figure 1E). These pathways, which are differentially regulated across hematopoietic, epithelial, and stromal cells, represent cell-intrinsic sex-based differences. For example, female cells have higher gene expression of metabolic, translation, and antigen presentation rates across all cell types. Male cells, in contrast, have increased gene expression of adipogenesis, proinflammatory signaling, and glucocorticoid signaling. Nine out the top 10 most differentially expressed pathways we identified for each sex were similarly sex-biased in human kidney^41^, suggesting that multiple cell types show consistent sex-biased enrichment of pathways linked to cellular metabolism, immune responses, and hormone signaling. Indeed, our data correspond closely with sex-based trends identified in human iPSC lines^42^ and other human organs^43^, indicating these pathways frequently show differences between male and female cells across many cell types.

In contrast to pathways with consistent sex-biased enrichment in thymic cell types, some pathways showed sex-biased enrichment in only a subset of cell types. For example, male T cells showed sex biased enrichment of integrin β1 and β3 pathways, a bias that was not found in other male cell subsets (Figure 1E). Likewise, only female T cells and epithelial cells showed sex-biased enrichment of slit and robo receptor signaling pathways. Our dataset also revealed a sex-specific shift with higher pathway enrichment for some cell types. In females, gene expression was consistent with higher cytokine signaling in T cells and hematopoietic cells, whereas in males, gene expression was consistent with higher cytokine signaling in epithelial and mesenchymal cells (Figure 1E). These data show that thymic cells from female and male donors show significant differences in gene expression. Given that higher levels of cytokine signaling has been previously shown to influence thymus and T cell biology^44,45^, our data suggests male and female T cells develop in different signaling environments and may respond differently to cytokine stimuli. This suggests that biological sex should be included as a variable in studies on immune cell and thymic organoid screening models as a critical parameter affecting cell growth, differentiation, and function.

Beyond expression-based differences, we quantified the cell type abundance within male and female tissue sections. The analysis revealed significant differences in the distribution of cortical and medullary cells between sexes. For example, when normalized to the total number of cells in each lobe, female thymus lobes contained significantly more DP cells (p = 0.011) and cTECs (p = 0.0023). In males, we found significantly more SPs (p = 4.2 x 10^-4^), CD3+ DPs (p = 9.9 x 10^-4^), activated mTECs (p = 0.0014), and VSMCs (p = 2.4 x 10^-6^) (Figure 1F). Given that thymus lobes with more DPs and cTECs would have a greater proportion of cells undergoing positive selection and more medullary cells would have more cells undergoing negative selection, these data suggest that a sex difference in cell type abundance may influence the resources directed towards specific stages of thymocyte selection. Alternatively, these results may suggest that male and female thymuses are developmentally asynchronous, with males exhibiting faster growth and involution kinetics, resulting in decreased cortical to medullary ratios even in early neonatal stages. Therefore, we focused further analyses on sequential developmental niches, including analysis of sex differences in key cell types and niche composition at each stage.

### JAG1 drives early thymic progenitor development towards thymic dendritic cells

We first focused on the cortico-medullary junction (CMJ) where cells home to the thymus (Figure 2A). This region recruits and supports early thymic progenitors (ETPs)^10^ and is composed of endothelial cells and pericytes expressing the Notch ligand JAG1 (Figure 2B,C). CITE-seq data demonstrated that the cell adhesion molecule used by ETPs to enter the thymus, CD62L, is quickly downregulated upon entrance through the vasculature at the CMJ (Sup. Figure 2A). However, recently immigrated CD62L+ double negative cells are frequently located in the subcapsular zone (Sup. Figure 2B). These data suggest that ETPs enter the thymus and rapidly migrate out to a subcapsular niche where DLL4, a more potent Notch ligand, is highly expressed on fibroblasts and subcapsular epithelial cells (Figure 2D, Sup. Figure 2C). The concentrated presence of JAG1 at their entry point, however, indicates that this is the first Notch ligand to which ETPs are exposed.

**Figure 2.**
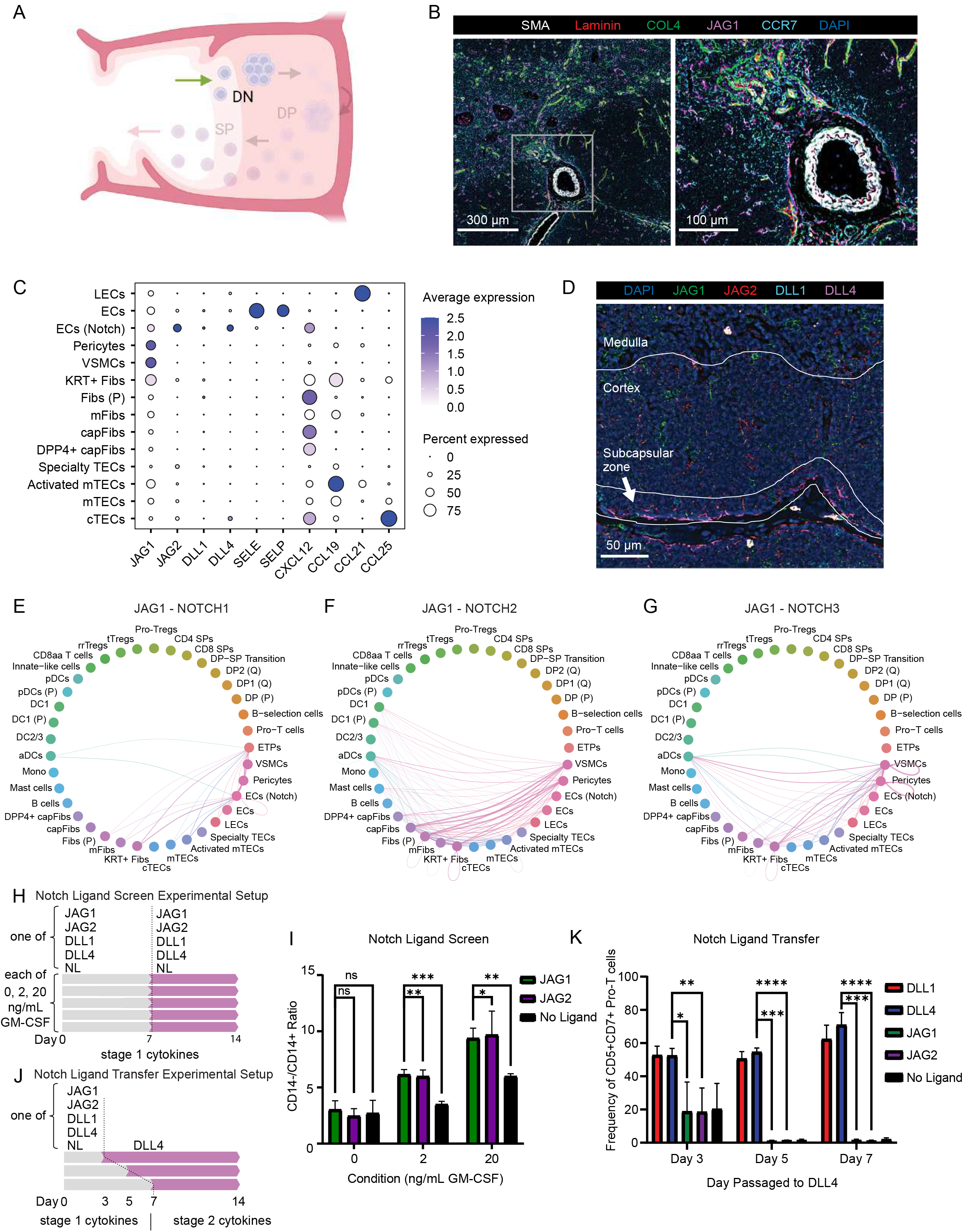
Thymic progenitors entering via the corticomedullary junction are exposed to a gradient of Notch ligands which influence lineage specification: A) Graphic depicting the ETP entry niche at the corticomedullary junction. B) Representative image of an arteriole niche at the corticomedullary junction. Vascular smooth muscle cells (white) make up arteriole walls and surround an inner lining of endothelial cells (red) which, along with local mesenchymal cells (green), express JAG1 Notch ligands (magenta). Early thymic progenitors expressing the chemokine receptor CCR7 (cyan) interact directly with JAG1 ligands after entering the thymus. C) Gene expression of important chemokines and Notch ligands by supporting thymic stromal and epithelial cells. D) Localization of the potent notch ligand DLL4 (magenta) to the subcapsular zone of the cortex. E) CellChat results of JAG1-NOTCH1, F) JAG1-NOTCH2, and G) JAG1-NOTCH3 interactions. H) Overview of *in vitro* Notch ligand screening experimental design. CD34^+^ hematopoietic stem and progenitor cells (HSPCs) were cultured on each of the four Notch ligands (JAG1, JAG2, DLL1, DLL4) or no ligands (NL) for 14 days. Each ligand condition was repeated in media containing 0, 2, or 20 ng/mL of GM-CSF. I) Bar plot showing effects of Notch ligand screens in the presence of varying levels of GM-CSF on the ratio of CD14^-^/CD14^+^ cells at the end of the 14 day differentiation protocol. All results shown are mean ± standard deviation from N=3 independent umbilical cord blood (UCB) donors. J) Overview of *in vitro* Notch ligand screening experimental design. CD34^+^ HSPCs were cultured on each of the four Notch ligands or no ligands (NL) and then transferred to DLL4 on day 3, 5 or 7. K) Bar plot showing results of Notch ligand transfer experiment on frequency of CD5^+^CD7^+^ Pro-T cells. All results shown are mean ± standard deviation from N=3 independent UCB donors.

Consistent with this notion, CellChat^46^ pathway analysis revealed that JAG1-NOTCH1 interactions between endothelial and perivascular cells are enriched with ETPs (Figure 2E), while JAG1-NOTCH2 and JAG1-NOTCH3 interactions are enriched with DC1, DC1 (P), DC2/3, and aDCs (Figure 2E-G). These data suggest that JAG1 could serve as a mechanism to induce commitment towards other hematopoietic lineages, such as pDCs, conventional dendritic cells (cDCs), or macrophages, some of which thymic development has been recently described in humans^10^. As JAG ligands have been shown to induce a lower level of Notch induction^47–50^, we hypothesized that early ETP contact could also act as a bridge to maintain T-lineage potential while cells migrate towards DLL4 in the subcapsular niche.

We first analysed the ability of the four Notch ligands present in the thymus to induce T-lineage commitment or alternative lineage development from cord-blood derived CD34^+^ hematopoietic stem and progenitor cells (HSPCs) in a defined, feeder-free culture system^44^ (Figure 2H). We included titrated concentrations of granulocyte-macrophage colony-stimulating factor (GM-CSF), which is produced by mast cells at the CMJ, and has been shown previously to support development of DCs^51^. We found that only DLL1 or DLL4 ligands could induce T-lineage commitment, whereas JAG ligands or no ligand controls supported development of myeloid cells (Sup. Figure 2D). Specifically, JAG ligands in the presence of GM-CSF skewed CD68^+^ DC development towards CD14^-^ DC1 cells compared to no ligand controls which skewed CD68^+^ DC development towards CD14^+^ DC2/3 cells (Figure 2I, Sup. Figure 2E).

Next, to test our hypothesis that Notch signals via JAG1 ligands could act as a bridge towards later DLL4 interactions, we analysed cells grown on JAG1 for 3 or 7 days prior to transfer to DLL4 (Figure 2J). We found that cells cultured on JAG ligands or no ligands for 3 days maintained reduced T-lineage commitment compared to DLL1 or DLL4 cells (p_JAG1_ = 0.033; p_JAG2_ = 0.017), whereas cells cultured on JAG ligands for longer than 3 days did not maintain T-lineage potential (Figure 2K).

Given that we observed gene expression consistent with a sex difference in signaling pathway activation, we next analysed the CMJ in males and females. We found significant sex differences in the contribution of different Notch ligands in the early development of ETPs (Sup. Figure 2F,G). Overall, our data suggest that JAG ligand interactions are more abundant and diverse in females. Specifically, we find JAG1- NOTCH1 interactions with ETPs only active in females and enriched DLL4 interactions in males (Sup. Figure 2H), which could lead to increased JAG1 interactions with ETPs in females as cells migrate towards the subcapsular zone.

Together, these data suggest that timely migration from the CMJ to DLL4 ligands at the subcapsular zone is critical for T-lineage commitment and that exposure to JAG ligands at the CMJ can guide alternative lineage development towards thymic-derived dendritic cells. Our data further reveal previously unrecognized sex-biased regulation of exposure to different Notch ligands.

### Analysis of the subcapsular zone reveals sex-based differences in fibroblast regulation of DP development and thymus growth

From the CMJ, ETPs migrate to the subcapsular zone via a CCL25-CCR9 chemokine gradient established by cTECs and directed to Pro-T cells, DP (P), and DP2 (Q) cells, but not DP1 (Q) cells (Figure 3A; Sup. Figure 3A). The subcapsular niche is defined by JAG1+ VCAM1+ DCs, cTECs, capsular fibroblasts, DPP4^+^ capsular fibroblasts, and a population of proliferating fibroblasts. These fibroblast populations secrete and maintain different ECM compositions which are tightly spatially regulated to support sequential thymocyte development (Figure 3B,C; Sup. Figure 3B,C).

**Figure 3.**
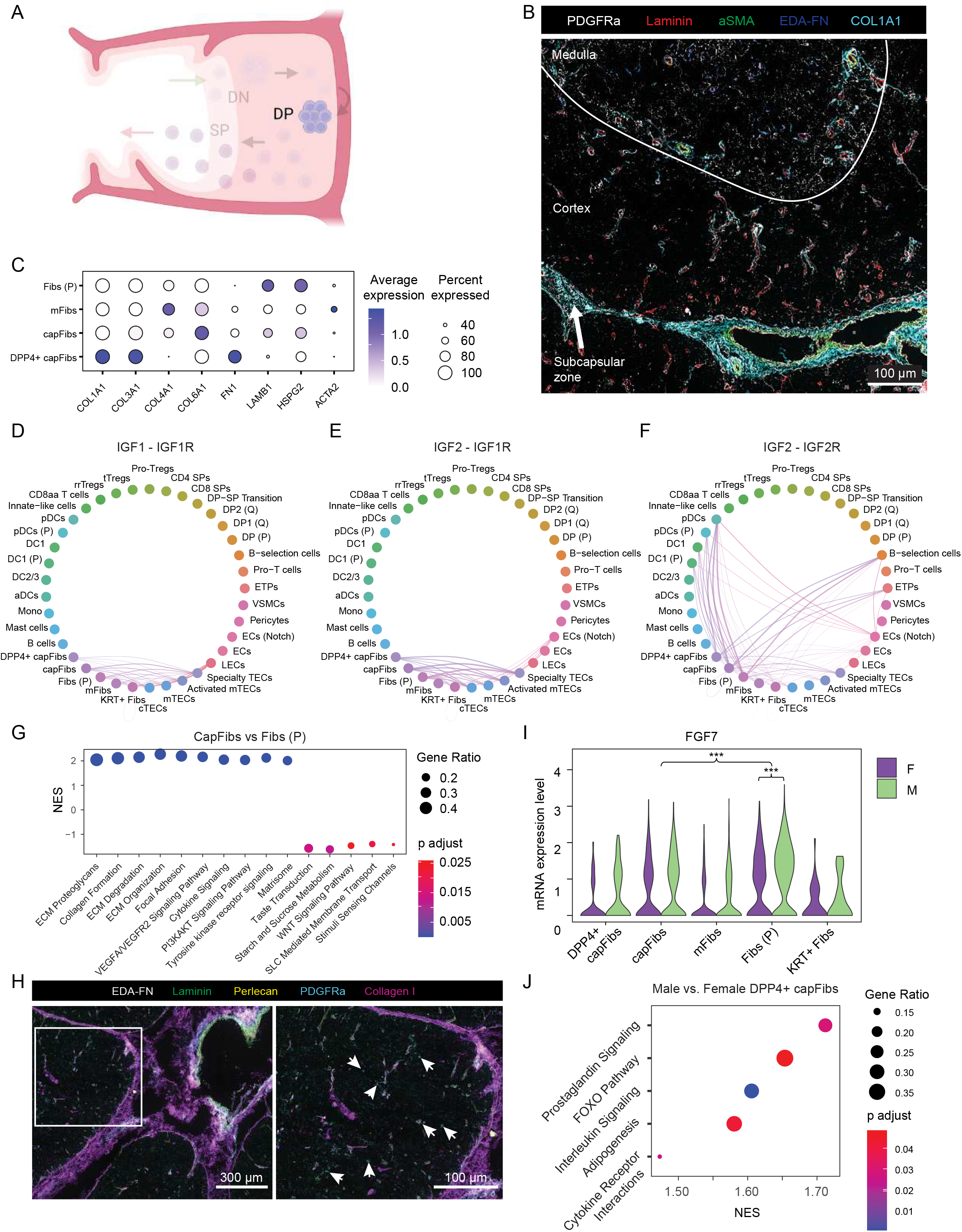
Fibroblasts in the subcapsular zone mediate thymus biology and T cell progenitor development: A) Graphic depicting the subcapsular zone. B) CODEX immunofluorescence image depicting the differential expression and spatial localization of various extracellular matrix (ECM) proteins within specific zones of the thymic medulla, cortex, and capsule. C) Gene expression data of different fibroblast populations captured via CITE-seq showing differential production of key ECM components. D) CellChat cell-cell interaction plots depicting interactions of IGF1-IGF1R, E) IGF2-IGF1R, and F) IGF2-IGF2R. G) GSEA dot plot of differential upregulated pathways between capFibs and Fibs (P). CapFibs have positive NES scores while Fibs (P) have negative NES scores. H) CODEX immunofluorescence image depicting a representative PGDRA+ ECM- cell niche within the early postnatal thymic cortex. Arrows point to PDGFRA+ ECM- cells located within the cortex. I) Violin plot depicting the differential expression of the key signaling molecule FGF7 by Fib (P) compared to other fibroblast populations, as well as upregulation by male vs. female Fibs (P); *** p<0.001. J) GSEA dot plot of differentially upregulated pathways in male vs. female DPP4+ capFibs. Male DPP4^+^ capFibs have positive normalized enrichment scores (NES) while female DPP4^+^ capFibs have negative NES scores.

We performed GSEA on our subcapsular fibroblast populations to determine functional differences and found that DPP4^+^ capsule fibroblasts were enriched in the HSP90 chaperone cycle for steroid hormone receptors (p_adjusted_ = 0.019; 21/52 pathway genes significantly upregulated) (Sup. Data 2). This suggests that DPP4^+^ capFibs are the primary fibroblast population which can respond to steroid hormones and implicates these cells as the orchestrators of sex hormone-based thymic involution. In contrast, capFibs were enriched for differentially expressed genes related to cytokine (IL33, p_adjusted_ = 1.50×10^-6^; IL34, p_adjusted_ = 3.56×10^-7^) and chemokine signaling (CCL2, p_adjusted_ = 5.10×10^-40^; CXCL3, p_adjusted_ = 0.020; CXCL12, p_adjusted_ = 1.78×10^-8^; CXCL14, p_adjusted_ = 3.63×10^-15^), indicating that these cells are important mediators of thymocyte migration and development. Furthermore, when compared to mFibs, capFibs were enriched in pathways related to insulin growth factor (IGF), which has been shown to regulate TEC proliferation and maintenance, as well as drive proliferation and suppress apoptosis in peripheral T cells^52^. Using CellChat we identified capFibs and Fibs (P) as major contributors to IGF signaling through predicted direct signaling to cTECs via IGF2-IGF1R and IGF1-IGF1R axes, and to ETPs and β-selection cells via an IGF2-IGF2R axis (Figure 3D-F).

Following this analysis, we next explored the role of our newly identified population of proliferating fibroblasts. We performed GSEA comparisons between our capFibs and Fibs (P) and found marked differences in signal transduction pathways, where capFibs resemble traditional fibroblasts upregulated in tyrosine kinase, angiogenesis, and ECM regulation and deposition pathways, whereas Fib (P) are upregulated in WNT signaling genes and in cell sensing pathways, including upregulation of genes involved in transient receptor potential (TRP) channels in the stimuli sensing channels pathway and taste receptors (TASRs) in the taste transduction pathway (Figure 3G; Sup. Data 2). Interestingly, within the cortex our spatial imaging data identified a unique population of ECM^-^ PDGFRa^+^ fibroblasts which lacked expression of EDA-FN, further indicating that Fibs (P) are not involved in fibrotic matrix deposition unlike capFibs (Figure 3H; Sup. Figure 3B). This population forms a network of PDGFRa^+^ cells throughout the cortex which does not overlap with the cTEC network, yet maintains cell-cell contact in specific niches, and contains localized populations of cells near cortical capillaries (Sup. Figure 3D).

Upon analysis, we found significant sex-specific differences in FGF signaling within this critical ECM^-^ fibroblast population. Although all fibroblast cells in the thymus produce a key cTEC growth factor, *FGF7*^53^, Fibs (P) express significantly higher *FGF7* than capFibs (p_adjusted_ = 1.46×10^-13^), and male Fibs (P) express significantly higher *FGF7* than female cells (p_adjusted_ = 4.86×10^-4^) (Figure 3I). Because *FGF7* has been implicated in regulating thymus size^53^, the sex bias in *FGF7* expression levels may contribute to the larger size of early postnatal male thymuses, which are larger than female thymuses in humans and primates^26^ (Sup. Figure 3E). Additionally, our CellChat results predict that male Fibs (P) have enhanced signaling through *FGF10*, which has been implicated in cTEC proliferation^53^, and only male VSMCs express *FGF18* (Sup. Figure 3F-H). Collectively, these results indicate that FGF signaling is a major regulator of TEC proliferation and maintenance and underlines its importance in in vitro models currently being developed to treat athymia and aging^54–56^ to support iPSC-derived cTECs.

Comparison of differentially expressed genes between male and female mesenchymal cells found a significant increase in gene expression of adipogenesis, prostaglandin, and FOXO signaling pathways in DPP4+ capFibs, a fibroblast population likely responsive to steroid hormone signaling (Figure 3J, Sup. Figure 3I). We also found a significant increase in the expression of *APOD*, a gene associated with downstream androgen, estrogen, progesterone, and glucocorticoid signaling^57,58^, across several male fibroblast populations (Fibs (P): p_adjusted_ = 1.12×10^-40^, capFibs: p_adjusted_ = 6.65×10^-4^, mFibs: p_adjusted_ = 2.46×10^-36^) (Sup. Figure 3I). Given that hormone signaling has been linked with thymic involution^29,59,60^, these data suggest thymic involution may already be initiating in early postnatal males

In sum, we identified three major roles for capsule fibroblasts within the subcapsular niche, including maintenance of tissue structure and organization through spatially defined ECM and chemokine signaling, direct regulation of cTEC maintenance and expansion, and a previously undescribed mechanism of direct coordination of T cell development through growth factors and cell-cell interactions.

### Human postnatal thymocytes self-select in the cortex to support positive selection of conventional αβT cells

Upon exiting the subcapsular zone, DP cells migrate into the inner cortex towards the medulla where they receive positive selection signals that guide T-lineage branching towards CD4 or CD8 SP cells (Figure 4A). For DP cells to transition towards the CD4 lineage, cells must receive TCR stimulation through MHC class II complexes, yet previous research in mouse has shown that transcriptional expression of MHC class I and II by DPs is specifically downregulated as cells transition to DP state^61,62^. This low transcriptional expression is hypothesized to prevent thymocyte-thymocyte self-selection during positive selection in the cortex, instead requiring DP cells to interact with cTECs to receive positive selection signals.

**Figure 4.**
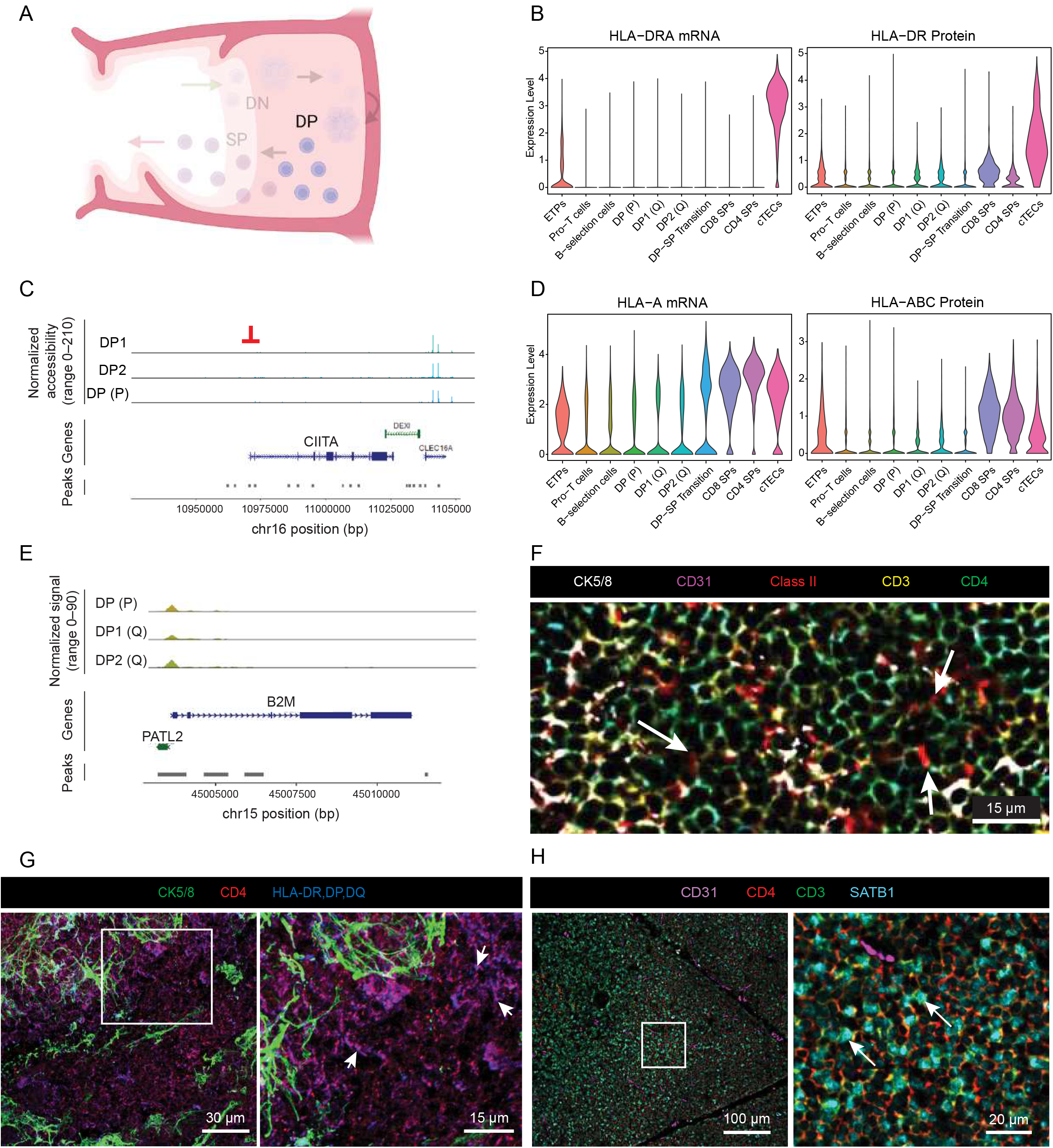
Class I and class II HLA interactions support thymocyte positive selection in the inner cortical zone: A) Graphic depicting the inner cortical zone. B) Violin plots depicting CITE-seq expression of HLA class II RNA and protein on developing thymocytes. cTECs are included for comparison of expression levels. C) Coverage plot of chromatin accessibility of the *CIITA* promoter in DP cells. D) Violin plots showing CITE-seq expression of HLA class I RNA and protein on developing thymocytes. cTECs are included for comparison of expression levels. E) Coverage plot of chromatin accessibility of the *CIITA* promoter in DP cells. F) CODEX immunofluorescence image depicting examples of class II^+^ (red) DP thymocytes (here shown as CD4^+^ (green)) interacting in the cortex. Arrows point to examples of class II at the junction of DP cells. G) Maximum projection confocal image showing lack of epithelial cells (Cytokeratin 5/8^+^; green) within specific niches of the cortex where class II^+^ (blue) DP cells (CD4^+^; red) reside. Arrows point to class II^+^ DP cells. H) CODEX immunofluorescence image depicting the inner cortical niche specialized for DP-SP transition, identified by SATB1^+^CD3^+^ DP (CD4^+^) cells. Arrows mark SATB1^+^CD3^+^ DP (CD4^+^) cells.

Analogous to the mouse literature, quiescent DP cells captured via CITE-seq do not express MHC class II transcripts and DP cells captured via ATAC-seq have closed CIITA promoters (Figure 4B,C). Despite the lack of class II mRNA, CITE-seq data demonstrated that thymocytes express low levels of MHC class II protein throughout development (Figure 4B). Additionally, in contrast to data in mouse, we observe constitutive class I mRNA expression which increased as cells transitioned towards SP cells (Figure 4D). These findings are consistent with ATAC-seq data demonstrating that the B2M promoter is open throughout thymocyte development (Figure 4E). We confirmed MHC expression via flow cytometry on postnatal thymocytes, finding that approximately 25% of DP cells are positive for both class I and II, and that over 65% of DP cells are class I^+^ (Sup. Figure 4A) Thus thymocyte self-selection within the cortex could support positive selection. In support of this notion, imaging data enabled us to identify locations within the cortex which did not contain epithelial, fibroblast, endothelial, or dendritic cells, but instead contained tightly packed DP cells expressing class II^+^ molecules concentrated at cell junctions (Figure 4F). We confirmed the absence of spindle-like cTEC projections in this niche via confocal imaging (Figure 4G). Finally, we quantified cell-cell interactions and identified a niche (Positive selection niche 1) consisting of class II^+^ DP cells and CD3^+^ DP cells, as well as a niche (Self-selection niche) containing mainly class II+ DPs (Figure 1D).

Next, we were particularly interested in identifying a niche which directs T-lineage commitment towards CD4 or CD8 SPs. We performed differential gene expression analysis on an early CITE-seq dataset lineage branch point to screen for markers to include in our deep imaging panel (Sup. Figure 4B). We found that SATB1 expression increased as DP cells transitioned towards SP cells (Sup. Figure 4C), and that compared to CD8 SP transition cells, CD4 SP transition cells had significantly higher expression of this master-regulator transcription factor^63^ (Sup. Figure 4D-E). Imaging analysis confirmed that increased expression of SATB1 coincides with CD3 upregulation, consistent with a role in late DP development and lineage branching (Figure 4H)^7^. Our neighbourhood analysis identified a niche enriched for mature CD3^+^ DPs in the inner cortex, suggesting that there either exists a niche specifically for late DP development and CD4 lineage transition, or that cells are pre-disposed to CD4 lineage development through their TCR and migrate as clonal populations after proliferation at the outer cortex.

Using image analysis to compare cortical niche organization between sexes, we found significant differences in how these niches are organized to support conventional T cell development, self-selection, and cross presentation. Females showed increased neighbourhood interactions between the cortical DC niche containing JAG1^+^ VCAM^+^ DCs and the mature DP niche containing CD3^+^ DPs, the positive selection niche 1 containing class II^+^ DP cells and CD3^+^ DP cells, and the positive selection niche 3 containing DCs and DPs (Sup. Figure 4F), and increased cell-cell interactions between cTECs and class II^+^ cells (Sup. Figure 4G,H). Conversely, males had increased cell-cell interactions between cTECs and CD3^+^ DPs (Sup. Figure 4G,H). These data suggest that the proportionally larger female cortex could increase cross presentation from DCs to cTECs, and possibly to class II+ DPs for self-selection, to facilitate greater use of this alternative mechanism for positive selection.

When taken together, our spatial multiomic analysis of the inner cortex identified several niches within the cortex supporting specific stages of DP development, including three conventional positive selection niches, a specialized niche for self-selection, and a mature DP niche thymocytes migrate through prior to entering the medulla.

### Spatial multiomics reveals key mechanisms regulating negative selection niches in the medulla

Mature DP cells entering the medulla transition towards CD4 or CD8 lineages and are subjected to an increased stimulatory environment specialized for negative selection (Figure 5A). Within the medulla, cells specialized for negative selection localize around keratinized structures called Hassall’s corpuscles^64^. These structures appear during late prenatal development and are abundant in human postnatal thymuses. Due to the rarity of these structures in mice^65^, their development, composition, and function have remained elusive. Here, we demonstrate that these structures can be divided into three major components: an external epithelial border of highly keratinized cells, an inner border cells expressing the prostaglandin-degrading enzyme 15-PGDH (*HPGD)*, and a central PDGFRa^+^ mass (Figure 5B). Hassall’s corpuscles are known to produce thymic stromal lymphopoietin (TSLP)^64^, an analog of the cytokine IL-7, which activates DCs to increase expression of class II molecules and the co-stimulatory molecules CD80 and CD86. Importantly, when we subclustered our stromal cell populations we found a new population of KRT^+^ fibroblasts which resemble cells undergoing epithelial-to-mesenchymal transition (EMT)^66^ (Sup. Figure 5A,B). In our CITE-seq data we identified TSLP^+^ and 15-PGDH^+^ cells in KRT^+^ Fibs, mFibs, mTECs, activated mTECs, and aDCs (Figure 5C), implicating these cell types as potential contributors to the function of Hassall’s corpuscles. Finally, given the inner layer of 15-PGDH^+^ cells we explored the role of prostaglandin signaling networks and regulation within the medulla. We found that DC1 and specialty TECs express high levels of PGE2, whereas DC2/3 cells and monocytes express the *PTGER2* and *PTGER4* receptors, and aDCs express the *PTGER3* receptor (Figure 5C). These data suggest prostaglandin signaling is a major regulator of DC activity in the medullary Hassall’s corpuscles.

**Figure 5.**
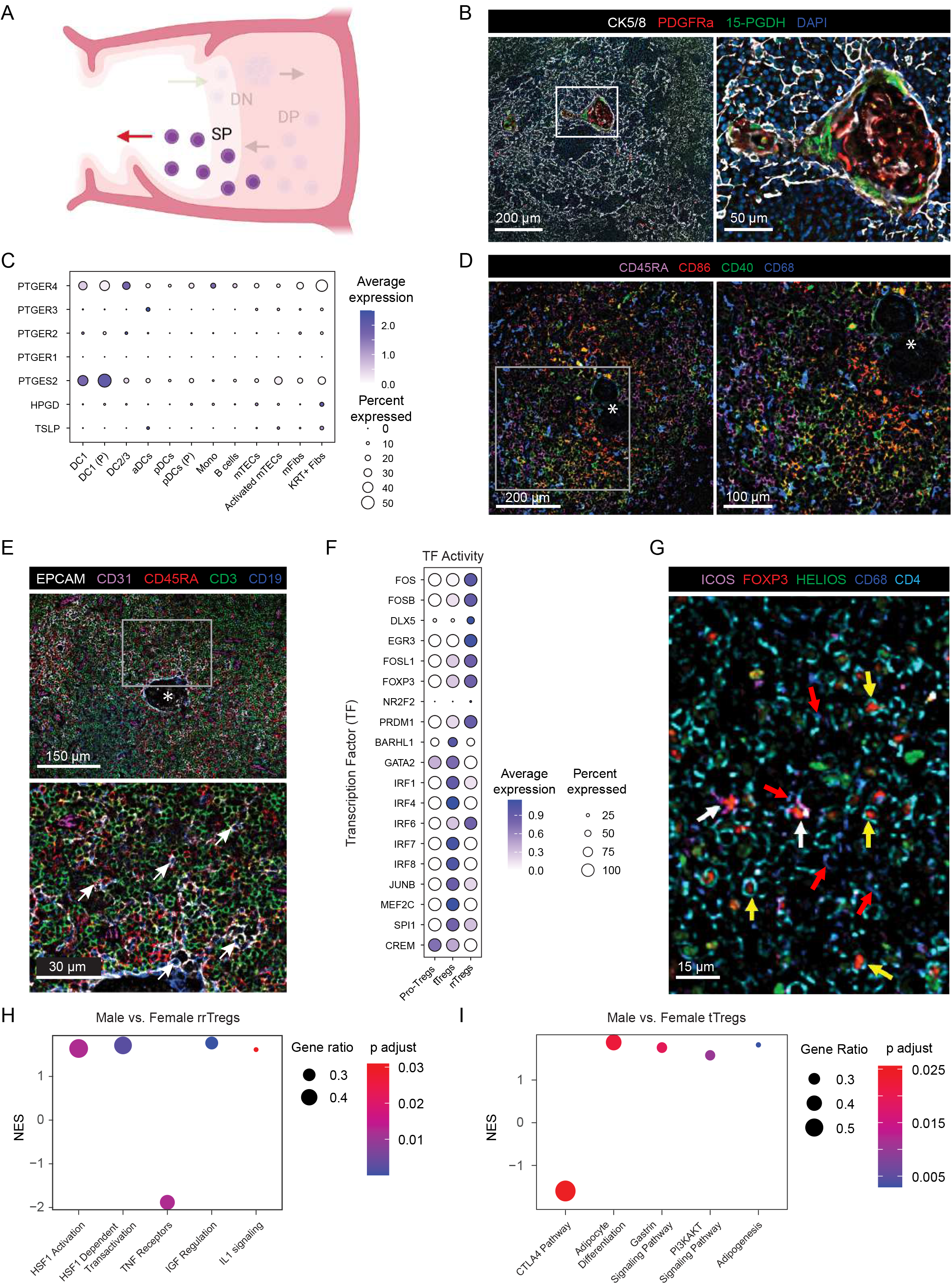
Hassall’s corpuscles represent scalable organizing centres for negative selection in the neonatal thymic medulla: A) Graphic depicting the thymic medulla. B) CODEX immunofluorescence images detailing three distinct components of human Hassall’s corpuscles. Cytokeratin 5/8 (white) creates an external border around an inner 15-PGDH+ (green) cell layer. PDGFRa (red) marks an inner core. C) Dot plot depicting gene expression of medullary-supporting stromal and epithelial cells contributing to key functional properties of Hassall’s corpuscles, as well as the hematopoietic cells responsive to Hassall’s corpuscle cells. D) Representative CODEX immunofluorescence images showing immune cells surrounding Hassall’s corpuscles. CD45RA (magenta) marks mature naïve SP cells, CD86 (red) marks antigen-presenting cells (APCs) which express CD40 (green), and CD68 (blue) marks key DC populations. Asterisk marks Hassall’s corpuscles. E) Representative CODEX immunofluorescence image showing B cells (CD45RA^+^, red; CD19^+^, blue) enveloped within mTECs (EPCAM^+^, white). Arrows point to examples of enveloped B cells. ￼F) Dot plot of SCENIC gene regulatory transcription factor results detailing transition from Pro-Tregs to tTregs, as well as the different transcription factor regulators governing rrTregs vs. tTregs. G) Representative CODEX immunofluorescence image showing rrTregs (CD4^+^, cyan; ICOS^+^, magenta) and tTregs (CD4^+^, cyan; ICOS^-^) located throughout the medulla and surrounded by CD68^+^ DCs (blue). White arrows point to ICOS^+^ rrTregs. Yellow arrows point to ICOS^-^ tTregs. Red arrows point to CD68^+^ DCs. H) GSEA dot plot showing differential expression of key regulatory pathways between male and female rrTregs. Male rrTregs have positive NES scores while female rrTregs have negative NES scores. I) GSEA dot plot showing differential expression of key regulatory pathways between male and female tTregs. Male tTregs have positive NES scores while female tTregs have negative NES scores.

CODEX imaging of medullary niches suggests that Hassall’s corpuscles act as sub-medullary organizational centers to segregate the inner human medulla into niches specialized for negative selection. We found that CD86^+^ APCs, a subset of which express the co-stimulatory ligand CD40, localize near Hassall’s corpuscles and are found in direct contact with CD45RA^+^ mature SP thymocytes (Figure 5D; Sup. Figure 5C). In addition, approximately 30% of medullary area is composed of B cells^67^. Again, we found that CD19^+^ B cells cluster into niches surrounding Hassall’s corpuscles (Sup. Figure 5D). These B cells are found in close contact with – and are often enveloped within – mTECs, indicating a potential role in cross presentation of antigens to epithelial cells (Figure 5E). Together these results suggest thymic B cells may comprise a major source of antigen generation that is passed to the network of epithelial and fibroblast cells for negative selection^67,68^, similar to their role in germinal centers of lymphoid organs. We quantified medullary neighbourhoods and identified six niches, including an mTEC maturation niche, a cross-presentation niche, and four niches specialized for negative selection which vary as to their location relative to Hassall’s corpuscles or the CMJ as well as the composition of APCs, epithelial, and T cells (Figure 1D; Sup. Figure 1C).

The negative selection niches organized around Hassall’s corpuscles play a key role in conventional T cell development, as well as the development of thymic Tregs^64^. To examine Treg development more closely, we sequenced samples enriched for CD25^+^ cells and examined gene and protein expression in subsets of Tregs. We found a population of CD25^hi^ Pro-Tregs which expressed canonical Treg markers *CTLA-4*, *TNFRSF1B* (TNFR2) and *TNFRSF4 (*OX40), positive/negative selection markers (*ITM2A, RANBP1, NCL, NME1, MIF, ATP5G1),* Treg developmental long non-coding RNA (*MIR155HG*)^69–72^, as well as other markers similarly described in mice (Sup. Figure 5E). Whereas Pro-Tregs expressed high levels of the pro apoptotic *BCL2L11*, mature tTreg subsets expressed the anti-apoptotic gene *BCL2*. Gene network reconstruction via SCENIC^73^ revealed transcription factor networks which are activated during pro-Treg to tTreg transition (Figure 5F).

In addition to newly developing tTregs, the thymus also contains mature, highly activated Tregs, labeled as rrTregs, which are thought to have recirculated from the periphery^74,75^. These rrTregs do not express *CCR7* or thymic egress markers (*KLF2, S1PR1*), but do express *IL1R2* (Sup Figure 5F), a receptor known to sequester the inflammatory cytokine IL-1β to reduce local concentrations^76^. CODEX imaging identified tTregs and rrTregs dispersed throughout the medulla, with rrTregs primarily adjacent to DCs marked by CD68 expression (Figure 5G). The potential of rrTregs to sequester inflammatory cytokines was supported by CellChat analysis, which showed interactions between DC2/3s and rrTregs through the IL- 1β - IL-1R2 axis (Sup. Figure 5F,G). rrTregs also exhibited a tissue resident Treg phenotype (*BATF^high^ CCR8^+^*) associated with wound healing and tissue regeneration function^77^, and indeed they expressed remodeling and tissue repair-related genes such as the matrix metalloproteinase enzymes (*MMP25* and *ADAM19*) (Sup. Figure H). In sum, these results suggest rrTregs may have an important anti inflammatory and tissue regenerating role within the thymic medulla.

Comparisons of male and female rrTregs revealed that male rrTregs had significantly higher expression of genes in IGF, HSF1, and IL1 signaling pathways (Figure 5H). Higher activity of the latter pathway suggests that rrTreg-mediated regulation of IL1R2-mediated anti-inflammatory feedback checkpoints is a more prominent mechanism in male tTreg development in early postnatal thymus. Of note, in activated mTECs GSEA revealed that males have significantly higher expression of inflammatory pathways such as CD40 and TNF than females, possibly resulting in higher rrTreg activity (Sup. Figure 5I).

Finally, we investigated thymic involution within the medulla in both males and females. As Tregs have been previously shown in mouse to contribute to thymic involution through a JAG1 associated mechanism^80^, we next explored sex-based differences in Treg gene expression. Using GSEA on Treg populations, we found that male tTregs showed significantly higher expression of genes associated with adipogenesis pathways (Figure 5I). Given the presence of cells undergoing EMT, our data underlies the aggressive timeline of thymic involution and suggests that sex-based differences in the decline of thymic function begins early in life.

Together, our detailed examination of the medulla shows that this zone is composed of several niches specialized for negative selection, cross-presentation, and maturation of mTECs which localize around Hassall’s corpuscles, and that these niches show sex bias in inflammatory pathways and thymic involution.

## Discussion

We performed spatial multiomics to construct a tissue atlas of niches guiding T cell development in human postnatal thymus. We employed these diverse datasets to characterize how key developmental niches drive lineage branch decisions, identify a new mechanism for conventional αβT cell development through self-selection, and identify new functions for recently described mesenchymal cell types governing thymus biology. Furthermore, we identified several sex-specific differences in thymus cell and niche biology. As T cell development is an inherently dynamic migratory process, knowledge of cell position in combination with proteomic, transcriptomic, and epigenomic sequencing data provides an invaluable set of data to predict niche-specific signaling cues directing specific stages of T cell development, as well as mechanisms responsible for maintaining tissue structure and contributing to thymic involution.

Our approach was to build an integrated spatial multiomic platform that would allow us to study the dynamic, spatially-regulated developmental processes in human thymic tissue. Here, we have chosen to highlight properties of niches which provided insight into specific questions of interest and were of fundamental importance to human T cell development. However, these findings describe only a subset of the data. We encourage the community to capitalize on the potential of this resource to provide insight into sex-specific differences and answer targeted niche-specific inquiries.

We describe a unique approach to sequencing data analysis where multidimensional spatial imaging serves as a benchmark for the location, ligand expression, and composition of key niches in T cell development. We then use niche composition to guide analysis of cell-cell interactions, allowing us to screen and identify ligand-receptor interactions which are physiologically relevant based on cell proximity in the tissue. In sum, this approach allows us to map epigenomic, transcriptomic, and proteomic data to distinct niches within the tissue at single cell resolution.

Previous thymus atlases built comprehensive transcriptional datasets to query T cell development from heterogenous ETPs to mature T cells, as well as describe the diversity of epithelial and stromal cells supporting T cell development^7,8,13,20^. We have expanded on these transcriptional datasets by enriching for and characterizing further diversity in Tregs and fibroblast cell types. Importantly, our atlas also includes surface proteome and epigenetic data for all sequenced cells and links single cell sequencing data to multidimensional spatial imaging and cell composition data for each sample. Finally, we included equal numbers of male and female age-matched thymus samples, enabling a unique opportunity to compare between sexes across platform modalities.

Our analysis of sex-matched human early postnatal thymus has revealed a highly plastic nature of thymus lobule organization and resource dedication, where each niche is responsive to sex and the corresponding differential developmental kinetics. This dynamic organization enables the diversity of developmental fates needed for T cell and hematopoietic cell development in the thymus. It also points to an underlying robustness in T cell development in that ultimately functional immune systems can arise in different manners. Major findings of sex-biased developmental thymic niches are outlined in Figure 6.

**Figure 6.** The human early postnatal thymus lobule is spatially organized into sex-biased niches to support stage-specific T cell development: A) In the cortico-medullary junction (niche 1) ETPs migrate into the thymus lobule. If the ETP migrates to the subcapsular zone and interacts with DLL1 or DLL4 ligands it can differentiate into a Pro-T cell. Alternatively, it can differentiate into a DC2/3 cell if it does not interact with Notch ligands, or differentiate into a DC1 cell if it interacts with JAG ligands. ETP interactions with JAG ligands are predicted to be greater in the female early postnatal thymus. B) In the subcapsular zone (niche 2) capFibs interact directly with cTECs and DP cells through secreted growth factors. DPP4+ capFibs and ECM^-^ Fibs also secrete key signaling molecules, with higher expression of signaling molecules observed in male cells. Higher FGF7 expression by male ECM^-^ Fibs causes increased proliferation of cTECs, which could account for the larger male early postnatal thymus. C) In the inner cortex (niche 3) DP cells can self-select off other DP cells that express class II HLA molecules. These self-selection interactions, as well as cross presentation of antigens from DCs to cTECs, are enriched in the early postnatal female thymus. D) In the inner medulla (niche 4) Hassall’s corpuscles (HC) act as organizing centers for medullary B cells, DC2/3, and DC1 cells. HCs play a role in prostaglandin signaling regulation, where DC1 cells produce prostaglandin (PTGES2) which acts on DC2/3 cells through the receptor PTGER4, and HCs express the prostaglandin degrading enzyme HPGD to decrease local concentration of PTGES2. HLA^high^ Activated mTECs in this niche have increased expression of inflammatory molecules CD40 and TNF. To counteract this increased inflammatory environment, male rrTregs express higher levels of IL1R2 to sequester the inflammatory cytokine IL1β produced by DC2/3 cells, inhibiting the costimulatory properties of DC2/3 cells.

In specific cases, such as the analysis of Notch ligands, we complemented our *in silico* approach with *in vitro* analysis. Our analysis suggests that JAG1 at the CMJ cannot support T-lineage commitment as cells migrate towards the subcapsular zone, but instead function to skew alternative lineage development at the CMJ towards a CD14^-^ DC1 subset (Figure 6). CD14 expression on DCs has been linked with increased inflammatory cytokine production^81^, suggesting that JAG ligands skew DC development towards non inflammatory DC phenotypes. Together these results highlight the importance of carefully tuned Notch signaling strength and timing during T cell and alternative lineage cell development in the thymus, as well as emphasizes the need for strict spatial control of different Notch ligands within sequential thymic developmental niches. Our observation of high JAG1 expression in the medulla and decreased expression of DLL4 on cTECs outside the subcapsular zone aligns with previous studies on human postnatal thymus^82^. However, unlike previous work our study did not identify diffuse expression of DLL4 or DLL1 on mTECs in sequencing or imaging data^82^. Additionally, our sequencing data identifies two distinct populations of endothelial cells marked by high expression of Notch ligands JAG2 and DLL4. Future work should characterize the roles of these endothelial cell types in directing different populations of cells upon entry and exit.

This study did not account for known subsets of ETPs which enter the thymus and could contribute to the differential commitment towards myeloid and T cell lineages^10,13,16^. However, these results demonstrating DC1 and DC2/3 development via exposure to specific Notch ligands in a thymus-like engineered niche do support recent literature which used computational trajectory analysis to demonstrate intrathymic generation of conventional dendritic cells subsets^10^. That study also suggested the possibility of intrathymic pDC development but could not find clear marker genes to define progenitor pDCs^10^. We find a similar phenomenon, where our progenitor-2 and pILC subsets express some, but not all, markers of thymic pDCs, limiting our analysis of the contribution of Notch signaling to the possible intrathymic development of pDCs.

In the subcapsular zone we characterize important roles of specialized fibroblast cells, where DPP4^+^ capsular fibroblasts respond to systemic hormone levels and capsular fibroblasts contribute to DP development indirectly through maintenance of cortical epithelial cells, as well as directly through immunoregulatory growth factors. DPP4^+^ capsular fibroblasts have been described in mouse in the thymic capsule^83^ and elsewhere as cells with progenitor and anti-fibrotic potential^84–87^. Here, we observe that DPP4 marks a subset of capsule fibroblasts which can respond to changes in systemic hormone levels. Thymic function and degeneration through thymic involution are known to be orchestrated through sex hormone levels^60,88–90^, implicating this DPP4^+^ capsular fibroblast population as a regulator of these processes and as a potential target for emerging strategies to address age-related thymic involution^91^. Additionally, whereas previously only medullary fibroblasts have been implicated in direct contribution to thymocyte development and selection in the medulla^83^, we demonstrate that capsule fibroblasts have the potential to support thymocyte development in the cortex through production of key growth factors (Figure 6). These cells represent a new source of cytokines and growth factors which can be mined for use in *in vitro* developmental systems. Finally, we demonstrate that ECM profiles of thymic fibroblasts are tightly regulated based on spatial localization within the tissue. Future work should characterize how tissue stiffness changes as cells migrate through developmental thymic niches to inform biomaterial strategies which support *in vitro* T cell development^92^.

Furthermore, using our integrated approach we identify a novel population of ECM^-^ cortical fibroblasts via multidimensional imaging and confirm the presence of ECM^-^ proliferating fibroblasts in our sequencing data which are upregulated in cell sensing pathways, such as TASRs and TRP channels. Interestingly, TASRs have been shown in non-olfactory systems to play a regulatory role where cells detect local soluble substances, such as glucose, and respond through release of hormones and other signaling molecules^93^. Similarly, TRP channels have been shown to play important roles in cell sensing, such as pheromone signaling, nociception, temperature sensation, and osmoregulation, as well as contribute to motile functions such as vasomotor control^94^. Given the proximity of these cells to vasculature in the cortex, these data suggest Fibs (P) play a critical role as regulatory cells by sensing changes in the thymic environment and responding to modulate thymic size (Figure 6). The lack of ECM production and the network-like structure of these cells resembles fibroblast reticular cells (FRCs) in the lymph node, which upon infection rapidly proliferate and instruct remodeling of the cortex^95^. Our age matched thymus samples were collected from early postnatal (4-5 months old) donors, which corresponds to the age in which T cell development is most active and the thymus must grow to accommodate lymphocyte production. We propose that this population of fibroblasts plays a similar role in expansion of the thymic cortex as FRCs during infection and signal primarily through FGF and IGF pathways to surrounding stromal and epithelial cells to orchestrate remodeling.

While the dogma in thymocyte positive selection suggests that DP cells actively downregulate class II RNA to prevent self-selection and to force DP cells to migrate and interact with cTECs^61,62^, several studies have suggested that T-lineage cells can select off each other to support CD4 T cell development^20,96–98^. Here, we describe an inner cortical niche where class II^+^ DP cells reside and have the potential to support positive selection via DP-DP self-selection interactions (Figure 6). Additionally, we show that upregulated SATB1 expression identifies mature DP cells in an inner cortical niche and the CD4 branch of their progeny, suggesting that it may be an early determinant of lineage specificity. Future work should investigate critical features of this niche, as well as the role of SATB1 in thymocyte development.

Within the medulla we detail a niche adjacent to Hassall’s corpuscles specialized for negative selection and describe an important role of recirculating Tregs in modulating the medullary inflammatory environment (Figure 6). The abundance of Hassall’s corpuscles in human, but not mouse, thymus and their proximity to negative selection niches suggests that these structures may have evolved to provide niche level organization within the larger human medulla or to provide more stringent spatial regulation of negative selection in species with a longer lifespan. Future research into the presence and organizational roles in other species could provide insight into the role of these structures in effectively scaling the thymic medulla while maintaining stringent negative selection.

By comparing male and female tissue we uncover significant sex differences in both T cell and thymus biology. Clues into the mechanisms underlying important differences in humans and model organisms have emerged from studies on male and female individuals at post-pubertal stages due to the known role of sex hormones in regulating thymic involution^27,60^. Further supporting the role of sex hormones, studies show androgen blocker treatment stimulated an increase in *FOXN1* expression, decreased the rate of thymic involution, and increased the rate of rejuvenation^29,30,59,89,91^. Decreased production of recent thymic emigrants and smaller overall size in older males versus females have also been reported^26,28^. With respect to cell and niche level changes, some studies describe decreased numbers of AIRE^+^ mTECs with age, potentially predisposing females who maintain greater thymic function later in life to autoimmune disease^29^, as well as less interlobular fat in young female thymus^26^, suggesting that differences in thymic involution kinetics begin pre-puberty. However, current literature has not addressed how transcript-level sex differences may underlie functional differences in thymic and immune function in humans. Our analysis of thymic cell gene expression uncovers that female-derived stromal, epithelial, and T cells have significant upregulation of metabolic, translation, and antigen presentation pathways whereas male cells have increased adipogenesis, proinflammatory signaling, and glucocorticoid signaling. These differences in cell metabolism align with current literature describing transcript-level sex differences in other organs^41–43^ and highlight the need for sex-based cell culture optimization to meet differential cell growth and differentiation requirements in *in vitro* T cell culture systems.

In addition to these changes common to other organs^40,41^, we identify meaningful thymus-specific differences which could significantly affect key processes in the development and training of thymocytes. Females have a larger proportion of cortical cells per lobule which aligns with known slower rates of thymic involution in females leading to a larger cortex/medulla ratio^26,27,59^. ETPs have enriched interactions with JAG1 ligands as they migrate away from the CMJ suggesting increased JAG1 ligand interactions could skew a greater proportion of ETP lineage commitment towards less inflammatory DC phenotypes (Figure 6). In the female cortex we observe increased interactions between cTECs and class II+ DPs and increased neighbourhood interactions between cortical DC niches and positive selection niches, suggesting thymocyte self-selection may play a larger role during positive selection (Figure 6). Conversely, in the female medulla we observe decreased activation of inflammatory pathways and less medullary cells. In sum these data, including a larger ratio of cortical to medullary cell types, suggest that females invest more resources in generating a larger repertoire of DP cells than in deleting autoreactive cells through negative selection, which could contribute to observed sex differences in the prevalence of autoimmune disease in females^99^.

In males, we observe enriched DLL4 interactions with ETPs, which aligns with previous data from sex hormone ablation studies demonstrating that androgen levels are positively correlated with DLL4 Notch ligands on cTECs^29^. Within the male cortex we observe increased interactions with mature CD3^+^ DPs and cTECs, suggesting that a smaller cortex could limit proliferation post β-selection to leave sufficient space for adequate positive selection. In the medulla, male activated mTECs exhibit a significant increase in inflammatory pathway markers and male Tregs exhibit higher inflammatory modulation and activate pathways involved in thymic involution^80^. Upregulation of inflammatory modulation by male rrTregs may also act as a regulatory mechanism to account for the higher common level of proinflammatory signaling in male cells (Figure 6). Alternatively, in the accelerated kinetics of the male thymus the tissue has transitioned towards an inflammatory milieu and medullary environment conducive to production of T cells associated with later developmental timepoints. Interestingly, post-pubertal males have more Tregs and less CD4 T cells than females, which could be a product of a more inflammatory environment in the medulla that skews CD4 development towards the Treg lineage^31^.

Furthermore, we advance knowledge of sex differences in thymus size control mechanisms. Within the fibroblast populations we find significant differences in the expression of key growth factors, such as FGF family members, which could contribute to the significant size difference in male and female thymuses at this age (Figure 6). These data align with and extend other known sex differences in growth factor expression, such as the sex biased expression of growth hormone and IGF-1 in regulating the size of different tissues^100,101^. Importantly, indicate that sex-specific differences exist in the maintenance and growth of thymus structure early in life, which could dramatically skew how our T cells develop, and suggests that interventions to prevent thymic involution must begin early in life. We also establish an early transition towards an environment conducive to adipogenesis in males. These data align with findings in model organisms, where young male rats exhibit higher rates of thymic involution^59^ and male early postnatal primates have a larger overall area of interlobular fat^26^. Together, these factors define two possible mechanisms that lead to a male-female difference in thymus size and involution kinetics. Future studies are needed to test how the sex differences at the transcript, niche, and organ level identified here contribute to differential production and quality of T cells and to resolve how sex differences in other complex organs contribute to known differences in the immune response.

Understanding the inherent differences in male and female cell immunology and the specific requirements of iPSC-derived cells from different donors is increasingly important as immunotherapies are adopted into the clinic^31^. Here, we identify sex-dependent differences in male and female T cell, hematopoietic, epithelial, and mesenchymal cell signaling pathways which could significantly affect how cells respond to specific signaling environments. These data emphasize that comparing male and female cell lines when modeling disease *in vitro* should be a critical component to accurately assess drug responses in different patient populations.

This work is limited by access to patient samples and the inability to conduct mechanistic experiments in the context of a whole animal. However, defined *in vitro* and organoid culture systems which strive to recreate the thymic microenvironment present powerful platforms to begin testing the roles of the cell type-specific and sex-specific signaling pathways described here to determine how they might contribute to increased incidence of autoimmune disorders in the female population and increased infection risk in males should be pursued. Furthermore, given the surprising differences in males and females at this early postnatal stage, future work should examine aged thymus to investigate how differential thymic involution kinetics, catalyzed at the cellular level, may translate to larger impacts on our immune system later in life. For example, how do age-related changes in thymic niche composition skew TCR diversity of recent thymic emigrants? Our spatial multiomic approach to analysing rare human samples should be applied to aged, diseased, and autoimmune thymuses to investigate how tissue niche biology changes under aberrant conditions.

## Acknowledgements

We thank the parents of thymus donors for their essential contribution to this research, as well as the surgical and cardiac clinic staff at the British Columbia Children’s Hospital who made this study possible; special thanks to Drs. Sanjiv Gandhi, Andrew Campbell and Al Aklabi, as well as Allison Jamieson, Melanie Ganshorn, Aliyah Hassan, Lyn Nguyen and Colleen Ring. We also thank Dr. Sabine Ivison for logistical support in tissue collection. We would like to acknowledge T. Stach, Y, Chung, S. Yu, and B. Zhao from the BRC-seq for technical support, M. Williams from the BRC Antibody Lab for technical support and custom reagents, and A. Johnson and J. Wong from the UBC FACS for technical support. We also thank Jen Ma for technical support with graphics. The cartoon figures were made with BioRender.

## Funding

Funding for this work was provided by Genome BC (SIP031) and Wellcome Leap Human Organs, Physiology, and Engineering (HOPE) program and the Canadian Institutes of Health Research (ICC-17644 to M.K.L) L.N.S is supported by a National Sciences and Engineering Research Council of Canada (NSERC) Canadian Graduate Scholarship – Doctoral (CGS-D) award and received a Michael Smith Foundation travel award to complete this work. K.S is supported by Wellcome Leap Inc (AWD-017859). E.A.F. was supported by an NSERC Undergrad Student Research Award. J.M.E. was supported by an NSERC CGS-D award and Zymeworks-Michael Smith Laboratories Fellowship in Advanced Protein Engineering. M.C.M was funded by a NSERC USRA and CBR summer award. Y.X.W. was supported by the Canadian Institutes of Health Research (CIHR) (MFE-152457) and an NIH K99 award (K99 NS120278). G.C.B is supported by the Raven Fellowship. R.D.J. is supported by the Michael Smith Foundation for Health Research (Trainee Award #RT-2021-1946). H.M.B. was supported by funding from the Baxter Foundation, the Li Ka Shing Foundation, and the National Institute of Health (R01 AG069858 and R01 AG075436). E.J.R. holds a CIHR Sex and Gender Science Chair in Genetics and is a Michael Smith Foundation for Health Research Scholar. M.K.L. is a Canada Research Chair in Engineered Immune Tolerance and receives a Scientist Salary Award from the BC Children’s Hospital Research Institute. F.M.V.R. is supported by CIHR (38395). P.W.Z. is a Canada Research Chair in Stem Cell Bioengineering.

## Author Contributions

L.N.S., M.K.L., F.M.V.R., and P.W.Z. designed the study. L.N.S., K.S., J.M.E., B.ZB.L., and R.D.J. conducted the experiments. L.N.S., K.S., E.A.F., Y.X.W., G.C.B., M.C.M., and E.J.R. analyzed the data. L.N.S., K.S., F.M.V.R., and P.W.Z. wrote the manuscript. All authors reviewed the manuscript.

## Declaration of Interests

L.N.S., J.M.E., F.M.V.R., and P.W.Z. are inventors of an invention disclosure relating to dendritic cell differentiation from cord blood. P.W.Z. is a cofounder of Notch Therapeutics. The authors declare no other competing interests.

## Supplemental Figure Legends

**Figure S1.**
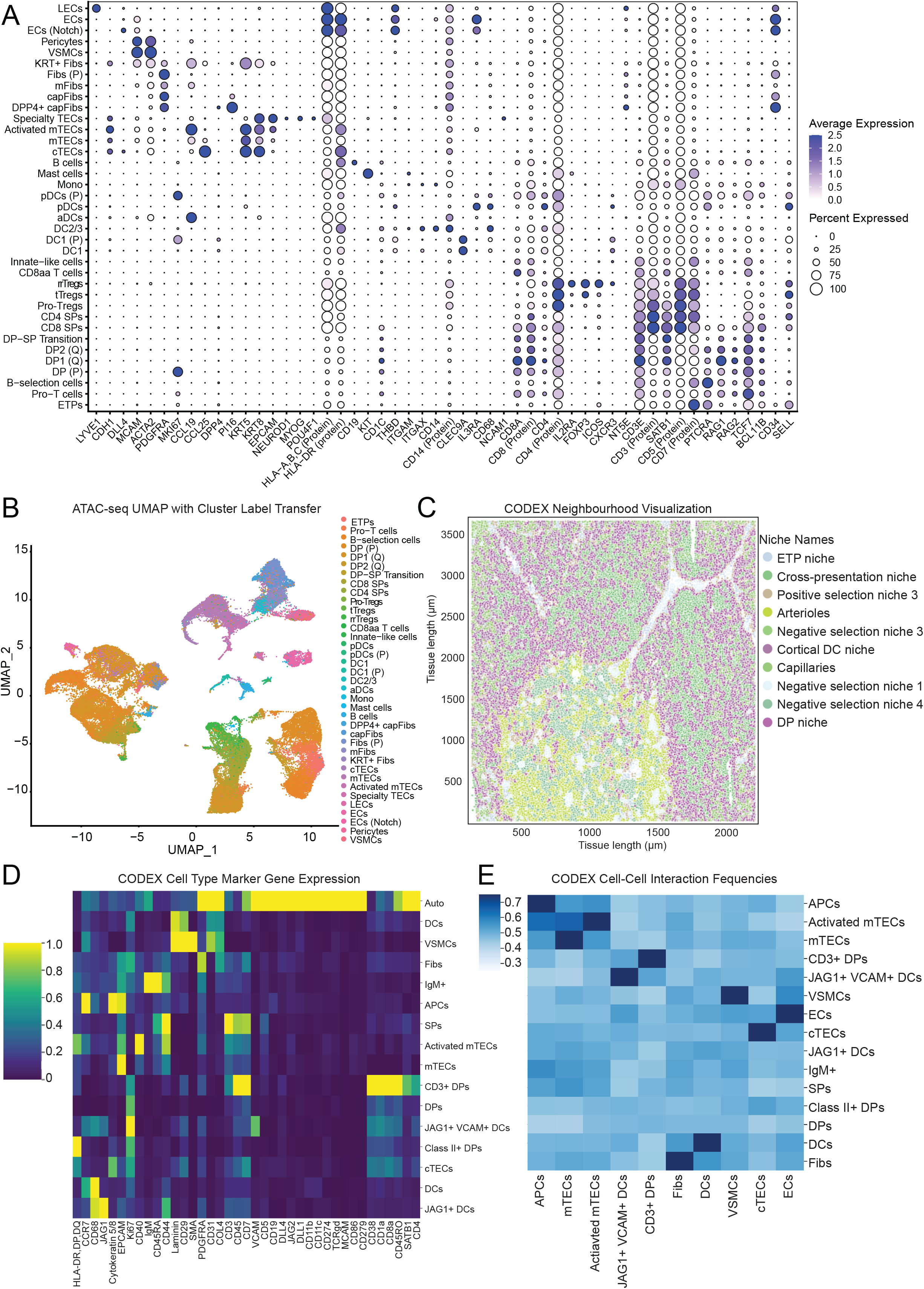
Spatial multi-omic analysis identifies specific cell types and tissue niches within tissue structure (related to Figure 1): A) Dot plot of key marker genes used to identify and align CITE-seq clusters with existing cell atlas datasets. B) ATAC-seq clusters identified by anchor-based label transfer from CITE-seq clusters. 34 out of 37 CITE-seq clusters are identified. C) CODEX neighbourhood visualization within a thymic lobule. D) Heat map of cell expression of phenotyping markers used to cluster cells identified via CODEX imaging. E) Heat map of individual cell-cell interaction frequencies.

**Figure S2.**
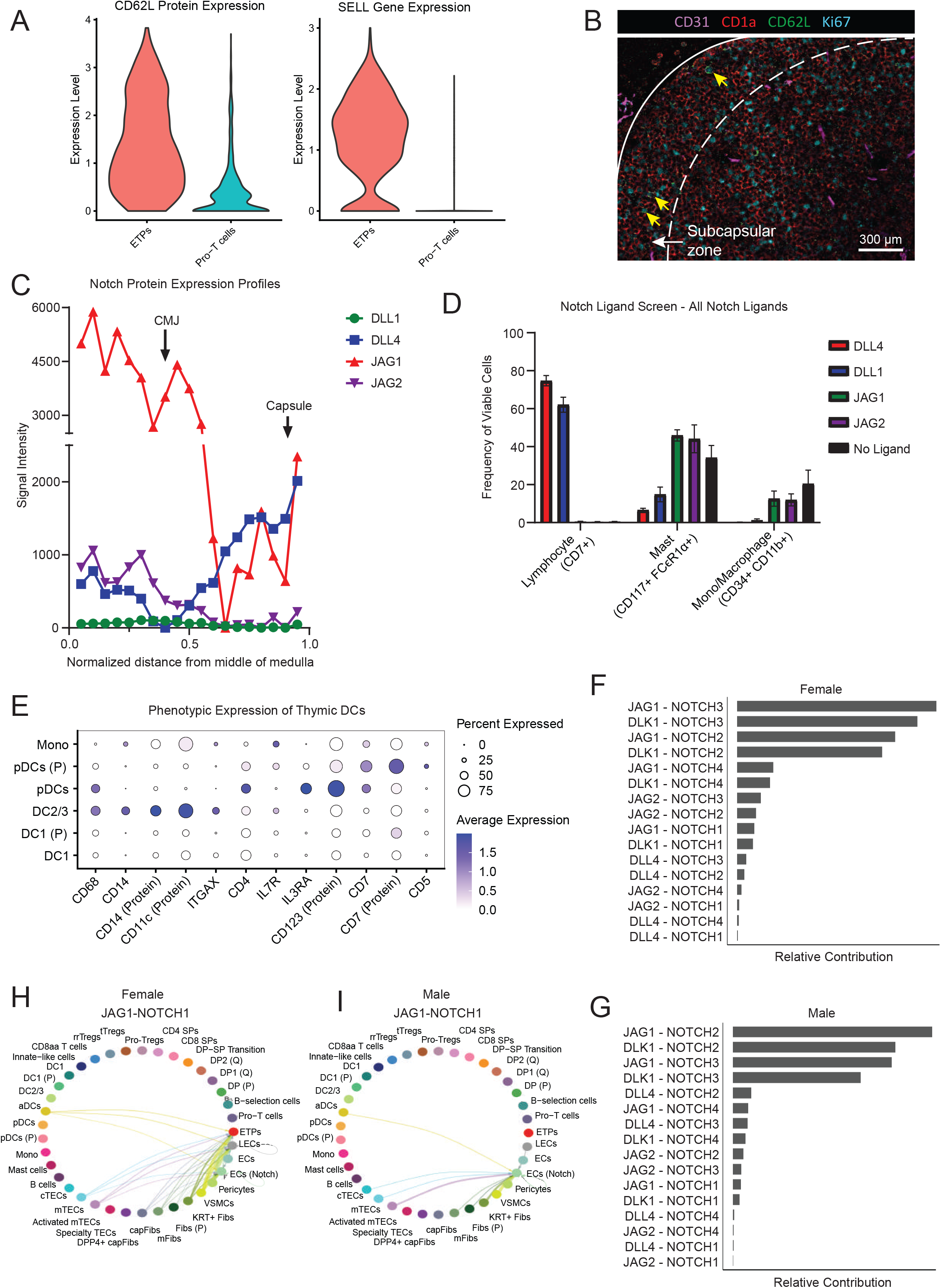
ETP specification at the corticomedullary junction is influenced by differential interactions with a gradient of Notch ligands (related to Figure 2): A) CD62L protein and gene expression dynamics during early thymocyte development captured via CITE-seq. B) CD62L+ ETPs localized at the subcapsular zone. Yellow arrows point to CD62L+ cells. The solid white line denotes the capsule and the dashed line denotes the boundary of the subcapsular zone. C) Notch ligand expression profiles across the thymus lobule. Ligand expression was measured from the middle of the medulla to the outer edge of the capsule. Three lines were drawn on each of three lobules from the center of the medulla to the edge of the cortex. D) Overall frequencies of various hematopoietic cell types which develop on different Notch ligands. Cell frequencies represent live cells in the 0 ng/mL GM-CSF condition. All results shown are mean ± standard deviation from N=3 independent UCB donors. E) Dot plot of key marker genes used to define DC CITE-seq clusters and align *in vitro* JAG ligand screening results with thymic-DC phenotypes. F) CellChat results of differential expression and utilization of Notch ligands by female and G) male cells. H) CellChat cell-cell interaction plot showing the differential interaction frequencies for the JAG1-NOTCH1 axes in female and I) male cells.

**Figure S3.**
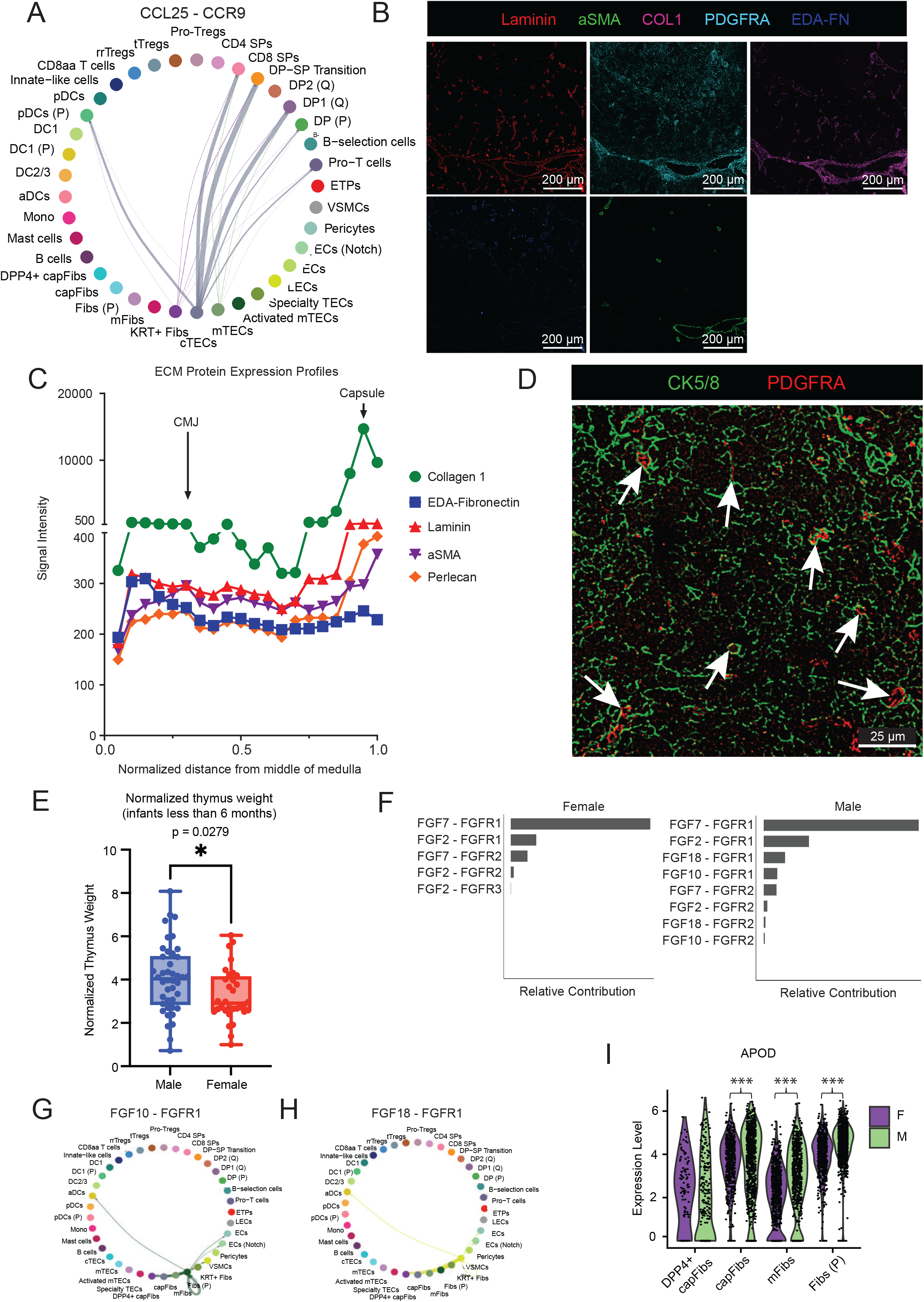
Cortical fibroblasts regulate tissue function through spatial regulation of tissue matrix proteins and cell growth factors (related to Figure 3): A) CellChat results of CCL25-CCR9 chemokine receptor ligand interactions directing migration to the subcapsular zone. B) CODEX immunofluorescence image of different ECM proteins found across the thymic lobule. Image shows strict spatial organization of various collagens and ECM proteins produced by different mesenchymal populations. C) Image quantification of various ECM proteins localized in distinct zones of the thymic lobule. ECM expression was measured from the middle of the medulla to the outer edge of the capsule. Three lines were drawn from the center of the medulla to the outer edge of the cortex for three different lobules. D) CODEX immunofluorescence image depicting distribution of fibroblasts (PDGFRa, red) and cTECs (CK5/8, green) in the cortex. Arrows point to fibroblasts in direct contact with cTECs. E) Box and whisker plot depicting the significant difference in normalized thymus weight of male and female donors. Due to limited patient data, thymus weights were normalized to Canadian weight standard for infants at each age^1^; * p < 0.05. F) CellChat results of differential expression and utilization of FGF signaling molecules by male and female cells. G) CellChat cell-cell interactions plot of FGF10-FGFR1 and H) FGF18-FGFR1 in male thymus. H) Violin plot showing the differential expression od downstream hormone signaling gene *APOD* in different fibroblast populations; *** p_adjusted_<0.001

**Figure S4.**
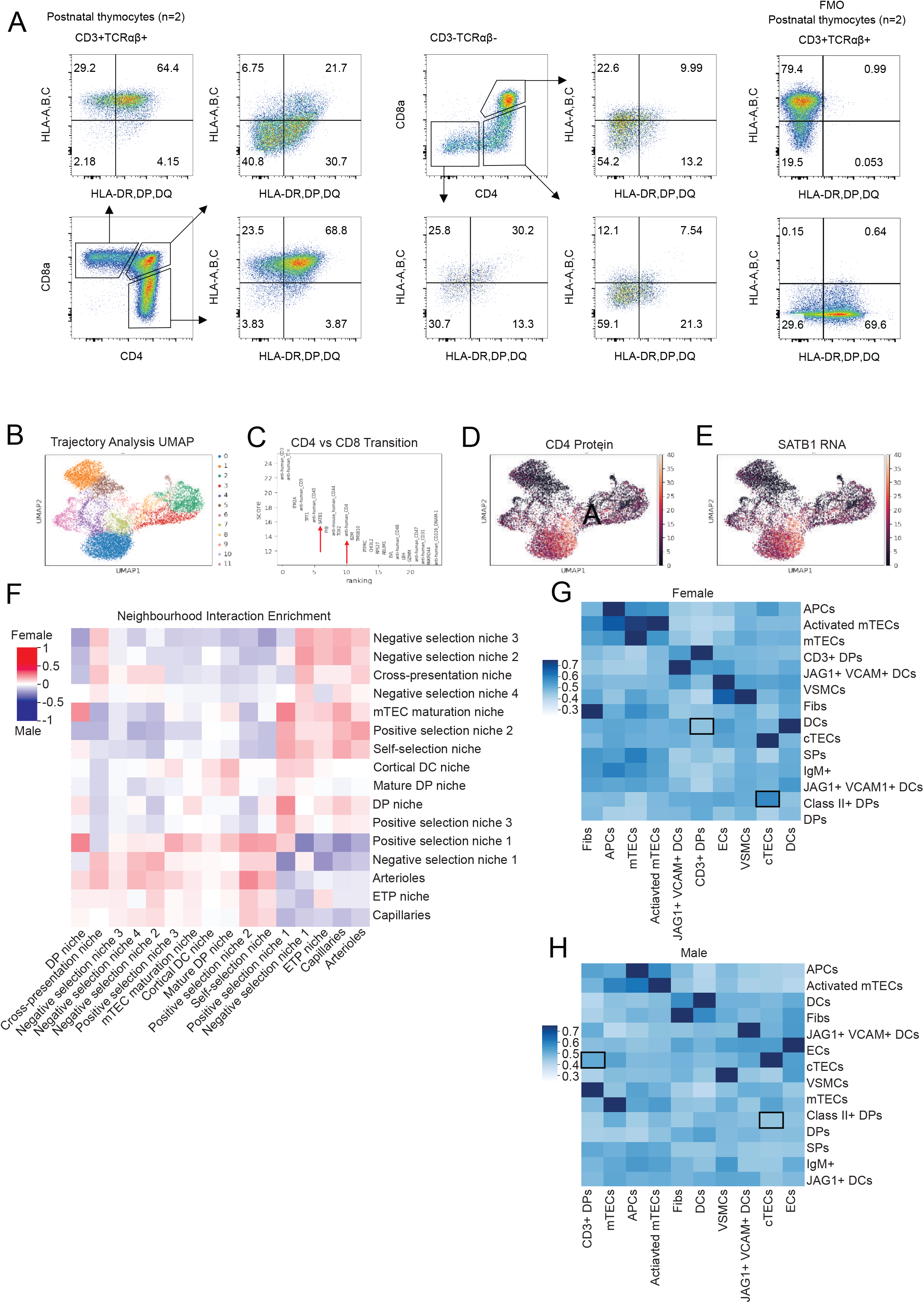
The inner cortical zone regulates positive selection and T-lineage specification through sex biased niche interactions (related to Figure 4): A) Flow cytometry data collected on bulk thymocytes (n=2). Lymphocytes were gated on FSC-A vs SSC-A and live cells, and then on CD3^+^TCRαβ^+^ co-expression. Cells were then gated on expression of CD4 and CD8a to identify different stages of thymocyte development. Expression of HLA class I and class II molecules on different subsets of T cells are shown. B) UMAP of first patient sequenced via CITE-seq and used to identify lineage branch point genes. C) Plot of differentially expressed genes in CD4 vs CD8 transition cells. D) UMAPs showing overlap in CD4 protein expression and E) *SATB1* gene expression. F) Neighbourhood interaction enrichment heatmap comparing female and male enrichment. The heatmap matrix was created by subtracting male enrichments from female enrichments. Red (upregulated) interactions are enriched in female thymus and blue (downregulated) interactions are enriched in male thymus. G) Female cell-cell interaction enrichment heatmap quantifying spatial imaging data. Black boxes highlight increased interactions between Class II+ DPs and cTECs as well as CD3+ DPs and DCs. H) Male cell-cell interaction enrichment heatmap quantifying spatial imaging data. Black boxes highlight increased interactions between Class II+ DPs and cTECs as well as CD3+ DPs and DCs.

**Figure S5.**
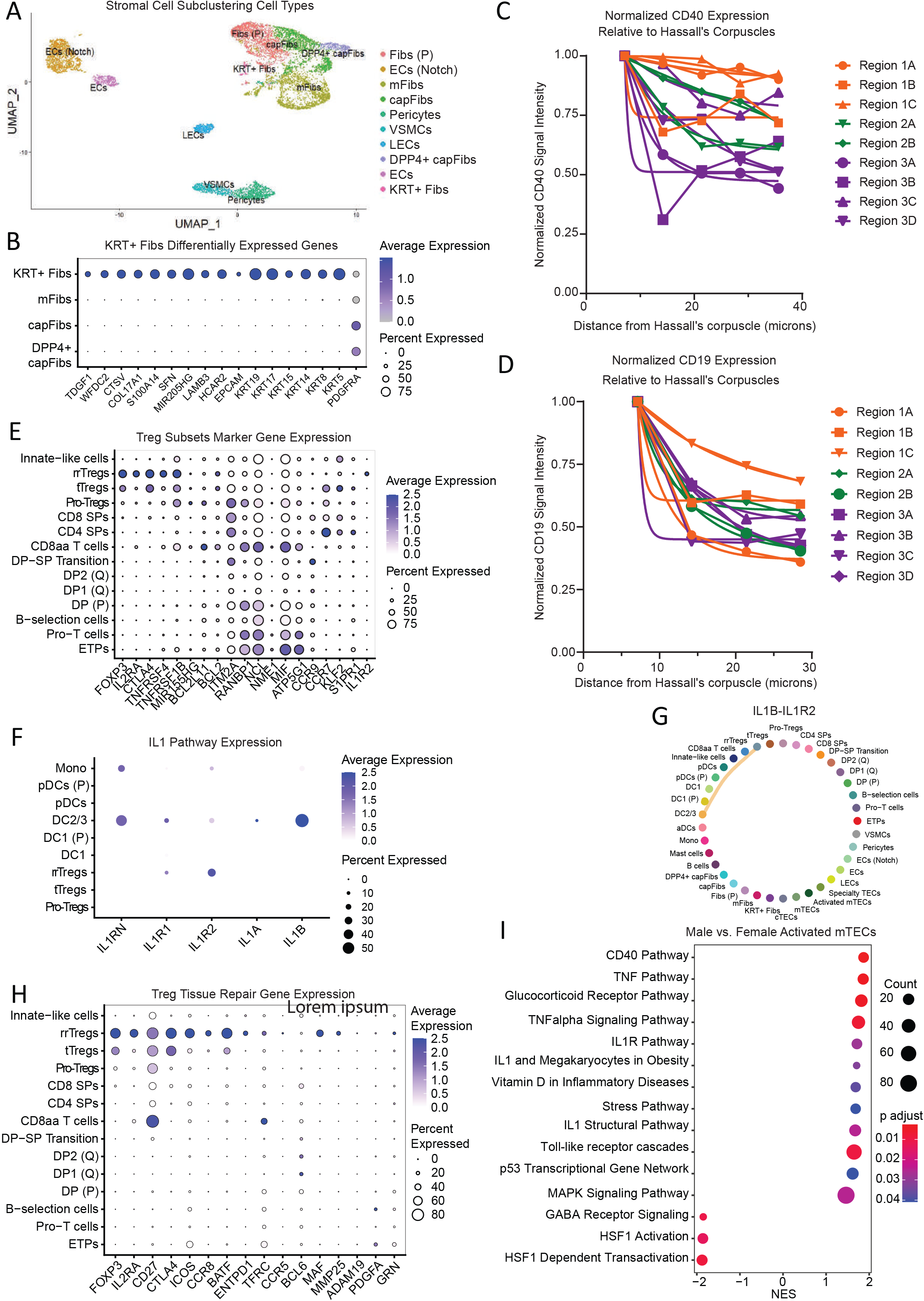
Regulation of negative selection niches are organized by Hassall’s corpuscles and regulated inflammatory pathway feedback loops (related to Figure 5): A) UMAP showing CITE-seq clusters identified after subclustering stromal cells. B) Dot plot of differentially upregulated genes in KRT^+^ Fibs compared to other fibroblast populations. C) Normalized CD40 expression relative to distance from Hassall’s corpuscles. A mask was drawn around each Hassall corpuscle and iteratively expanded, calculating mean CD40 expression intensity in the new mask area ring for each iteration. D) Normalized CD19 expression relative to distance from Hassall’s corpuscles. A mask was drawn around each Hassall corpuscle and iteratively expanded, calculating mean CD19 expression intensity in the new mask area ring for each iteration. E) Dot plot of key marker genes used to identify 3 Treg subsets. F) Dot plot showing differential expression of IL-1 pathway ligands and receptors across hematopoietic and Treg cells captured in early human postnatal thymus via CITE-seq. G) CellChat cell-cell interaction plot indicating an active IL1B-IL1R1 signalling axis between DC2/3s and rrTregs. H) Dot plot of key tissue regeneration and repair genes by rrTregs. I) GSEA dot plot showing differential expression of key regulatory pathways between male and female activated mTECs. Male activated mTECs have positive NES scores while female activated mTECs have negative NES scores.

## Methods

### Contact for reagent and resource sharing

Further information and requests for reagents and resources should be directed to and will be fulfilled by the lead contacts, Fabio M. Rossi and Peter Zandstra.

### Data and code availability

Deep imaging computational analysis and CODEX images have been deposited on Zenodo and are publicly available as of the date of publication. Due to the data size of raw CODEX datasets, down sampled versions of processed CODEX images are available (link). Unprocessed or full resolution images are available from the lead contact upon request. Codes for data analysis have been deposited at (https://gitlab.com/stemcellbioengineering). Any additional information required to reanalyze the data reported in this paper is available from the lead contact upon request.

### Study approval

Human research was approved by the University of British Columbia Research Ethics Board (H17-01490 and H18-02553).

### Experimental model and subject details

All tissue samples used for this study were obtained with written informed consent from parents/guardians of participants. Thymus tissue was collected during infant cardiac surgery at British Columbia Children’s Hospital and processed immediately upon retrieval. Tissue was collected in ImmunoCult-XF T cell Expansion Medium (immunoCult-XF; STEMCELL Technologies, Vancouver, BC, Canada) with 1% penicillin/streptomycin (P/S; Thermo Fisher Scientific) and processed in RPMI medium (Thermo Fisher Scientific, Waltham, MA, USA) with 10% fetal bovine serum (FBS). Upon collection the tissue was sectioned into 1cm^3^ pieces in RPMI + 10% FBS. One piece was used for CD45^-^ cell enrichment, one piece (∼3g) was used for CD25^+^CD8^-^ Treg enrichment using a custom thymic Treg selection kit (STEMCELL Technologies)^102^, two pieces were used for thymocyte isolation and cryopreservation, and the remaining pieces were processed for CODEX imaging.

### Pre-sequencing cell enrichment

The thymic tissue dissociation protocol detailed by Gustafson et al. was used to enrich for CD45^-^ cells^9^. Briefly, a 1 cm^3^ tissue section was crushed using a syringe in a small petri dish. Released thymocytes were collected in RPMI + 10% FBS, spun at 500xg for 5 min at 4°C, and kept on ice. The remainder of the thymic tissue was transferred to the lid of a 50 mL conical tube lid and minced using sterile surgical scissors. 8 mL of warm dissociation cocktail (2 mg/mL Stemxyme (Cedarlane Labs, Burlington, ON, Canada), 6.3 U/mL DNaseI (Sigma-Aldrich; Milwaukee, WI, USA), 1.5% BSA (w/v) (Wisent Bioproducts, Saint-Jean-Baptiste, QC, Canada) in RMPI) was added to the 50 mL tube, inverted 5 times to release thymic tissue from the lid, placed on a horizontal rocker, and incubated at 37°C for 30 min. After 30 minutes the supernatant was removed, filtered over a 50 μm strainer, and stored on ice (T3). An additional 8 mL of dissociation cocktail was added and the tissue was incubated for an additional 30 mins. After the second incubation the supernatant was removed, filtered, and stored on ice (T2). For the final 30 min incubation 8 mL of dissociation cocktail followed by 2 mL of 0.25% trypsin was added to the tissue fragments. After the final incubation the remaining cells were collected and filtered over a 70 μm strainer. 3 mL of FBS was added to the 50 mL falcon tube containing the final digestion media to break the trypsin reaction (T3). The three 50 mL collection tubes were spun at 500xg for 5 min at 4°C. Cells were resuspended in 40 mL of Hank’s Balanced Salt Solution (HBSS; Thermo Fisher Scientific, Waltham, MA, USA) with 2% FBS (HF), filtered through a 40 μm cell strainer, spun at 500xg for 5min at 4°C. Cells were resuspended in 4 mL of red blood cell lysis buffer (Millipore Sigma, Oakville, ON, Canada), incubated for 5 min at RT, diluted to a total volume of 40 mL with HF, and spun at 500xg for 5 min at 4°C. Cells were resuspended in 10 mL of HF and counted. The first incubation (T1) cells were pelleted (500xg, 5min, 4°C) and stored on ice for later use.

T2 and T3 cells were then depleted for CD45^+^ cells using the Miltenyi LD column prep kit (Miltenyi Biotec Inc., Charlestown, MA, USA). Briefly, T2 cells were added to T3 to reach a final cell number of 5×10^8^ cells. Cells were centrifuged at 300xg for 10min at 4°C and resuspended in 80 μL of HF buffer per 10^7^ cells plus 5 μL of CD45 beads per 10^7^ cells. The suspension was mixed well via gentle pipetting and incubated for 15 min at 4°C. Cells were washed by adding 1mL of HF buffer per 10^7^ cells and centrifuged at 300xg for 10 min at 4°C. During this incubation the LD column was prepared and washed with 2mL of HF buffer.

Pelleted cells were resuspended in 2.5 mL of HF and applied to the column. The column was washed with 2×1 mL buffer and all unlabeled flow through cells were collected as the negative fraction. The positive fraction was collected by passing 5 mL HF through the column using the plunger. Cells from each were counted prior to proceeding to antibody staining for cell sorting.

Both fractions were resuspended in Fc buffer (HF + 1:1000 human Fc block (BD Biosciences, Missisauga, ON, Canada)) to a final concentration of 0.5 M cells/mL. CD45-PE antibody (Biolegend) was added at 1:200 and the cell suspension was incubated for 30min on ice in the dark. After incubation cells were spun at 500xg for 5min at 4°C, washed with 5 mL of HF, and spun again. Cells were resuspended to a final concentration of 0.2 M cells/mL, stained with zombie UV live/dead stain (1:500; Biolegend), and stored on ice while transferring to the cell sorting facility. The CD45- fraction was sorted using an MOFLO Astrios EQ (Beckman Coulter) or BD Influx instrument into HBSS + 4% FBS until over 500,000 CD45^-^ cells were collected. The CD45^+^ fraction was then sorted to collect 500,000 live CD45^+^ cells. These cell suspensions were pelleted, counted, and kept on ice prior to processing for CITE-seq or ATAC-seq. Finally, to enrich Treg cells, we performed a CD25 positive selection on the bulk thymocytes, followed by CD8 depletion.

### CITE-seq sample preparation

The Biolegend TotalSeq-C CITE-seq antibody preparation and 10X Genomics CITE-seq cell staining protocols were followed. Briefly, prior to the end of the CD45^-^ sort one vial of TotalSeq-C Human Universal Antibody Cocktail v1.0 (Biolegend, San Diego, CA, USA) was equilibrated to room temperature for 5min and then spun at 10,000g for 30 seconds. The lyophilized panel was resuspended in 27.5uL of HBSS + 4% FBS, vortexed for 10 seconds, and then incubated at room temperature for 5min. The vial was vortexed again for 10 seconds and then spun at 10,000g for 30 seconds. The entire volume was transferred to a low protein binding PCR tube and then centrifuged at 14,000g for 10min at 4°C.

CD45^-^, CD45^+^, and CD25^+^ cell fractions were combined in a 2:2:1 ratio to a total cell count of 250,000 cells in a 12 x 75mm tube. Cells were spun at 500xg for 5 min at 4°C and resuspended directly in 12.5 uL of HBSS + 4% FBS. 12.5 uL of the antibody staining cocktail was added and the cells were incubated for 30min at 4°C. Cells were washed three times with 3.5mL HBSS + 4% FBS and resuspended in a final volume of 55 uL. Cells were counted, stored on ice, and given immediately to the sequencing facility for 10X Genomics 5’ library preparation and sequencing.

### ATAC-seq sample preparation

The 10X Genomics Nuclei Isolation from Mouse Brain Tissue for Single Cell ATAC Sequencing was followed. Briefly, CD45^-^, CD45^+^, and CD25^+^ cell fractions were combined in a 2:2:1 ratio to a total cell count of 250,000 cells in a 1.5mL Eppendorf tube. Cells were spun at 500g for 5min at 4°C. The supernatant was removed and 100uL of chilled 0.1X lysis buffer was added and mixed by gently pipetting 5x. Cell suspension was incubated for 5min on ice. Immediately after 5min 1mL of chilled wash buffer was added and the solution was gently pipet mixed 5x. Nuclei were centrifuged at 500g for 5min at 4°C, the supernatant was removed, and the nuclei were resuspended in diluter nuclei buffer to a final concentration of 5000 nuclei/uL. Nuclei concentration and acceptable morphology was confirmed via hemocytometer cell counting and brightfield microscopy, and a cell lysis percentage over 95% was confirmed using a Countess II FL Automated Cell Counter (Thermo Fisher Scientific). Nuclei were kept on ice and given immediately to the BRC sequencing core for single cell ATAC-seq library prep and sequencing.

The generation of single cell indexed libraries was performed by the Biomedical Research Center Next Generation Sequencing Core using the 10X Genomics chromium controller platform and the Chromium Single Cell 5’ Library and Gel Bead Kit v1.1 and Chromium Single Cell ATAC Kit v2 reagents. Briefly, around 10,000 cells were targeted per sample and deep sequencing was run on the NextSeq 2000 using the P3-100 cycle kit. After run completion, the Binary base call (bcl) files were converted to fastq format using the illumine bcl2fastq2 software, and data were received for further analysis.

### Notch ligand *in vitro* screening

Recombinant human JAG1-Fc, JAG2-Fc, DLL1-Fc (R&D Systems), or DLL4-Fc (Sino Biological) were diluted to 4 µg/ml in PBS along with 1 µg/ml VCAM-1-Fc (R&D Systems). To coat, 50 µl of each solution was added per well of a 96-well flat-bottom plate and incubated overnight at 4°C. Immediately before use, wells were washed once with 50 µl PBS before cells were seeded.

Umbilical cord blood was collected from consenting donors at BC Children’s Hospital, Vancouver, Canada in accordance with institutional research ethics. Mononuclear cells were isolated by density gradient centrifugation using Lymphoprep (Stemcell Technologies) and CD34^+^ HSPCs were enriched using the EasySep Human CD34^+^ Positive Selection kit (Stemcell Technologies) according to the manufacturer’s instructions. Isolated cells were cryopreserved in FBS containing 10% DMSO and stored in vapor phase nitrogen prior to use. Cells were thawed in 37°C water bath and 5x volume of 37°C Hanks Balanced Salt Solution with 0.5% BSA was added dropwise. Cells were centrifuged for 7 minutes at 250g, the supernatant aspirated, and resuspended in culture medium.

Cells were cultured in Iscove’s Modified Eagle Medium (IMDM) supplemented with 20% serum substitute (BIT 9500; Stemcell Technologies), 1 µg/ml low density lipoprotein (Stemcell Technologies), 60 µM ascorbic acid (Sigma), 24 µM 2-mercaptoethanol (Sigma), and 1% penicillin-streptomycin (Invitrogen). For experiments measuring T-lineage potential, cells were cultured in stage 1 cytokine concentrations from day 0-7 and stage 2 from day 7-14 (Sup. Table 5). Experiments investigating monocyte/macrophages and dendritic cells used stage 1 cytokines throughout the entire 14 days. All cytokines were from R&D Systems.

To begin experiments, 2000 cells per well were seeded on DLL1-Fc or DLL4-Fc, and 4000 cells per well on JAG1-Fc, JAG2-Fc, or wells without any Notch ligand. For experiments measuring T-lineage potential, cells were passaged to plates freshly coated with DLL4-Fc on day 3 or 7. For experiments measuring monocyte/macrophage and dendritic cells, cells were passaged to the same Notch ligand on day 7.

Cells were passaged in 100 µl of medium per well and an additional 100 µl was added 3-4 days after passaging, unless the cells were passaged (as on day 3), in which case the medium was completely changed, and the cells were seeded in 100 µl of medium. Gentle pipetting was used to passage cells, followed by rinsing wells with 30-50 µl of PBS + 0.5% BSA + 2 mM EDTA to remove any remaining adherent cells.

Cells were passaged, rinsed once with PBS, then stained for viability using Zombie-UV (Biolegend) diluted 1:500 in PBS. Afterwards, cells were incubated for 5 minutes with 1:100 FC block (BD Biosciences) in Hanks Balanced Salt Solution with 2% FBS (HF). For analysis of antibody panels containing CD68, cells were fixed and permeabilized using the Cytofix/Cytoperm kit (BD Biosciences) according to the manufacturer’s instructions. Surface and intracellular antibodies were stained together for 45 minutes on ice, then cells were rinsed twice with permeabilization buffer and resuspended in HF for analysis. For the antibody panels containing only surface markers, cells were stained in Brilliant Violet Buffer (BD Biosciences) for 10 minutes on ice, rinsed once with HF, then resuspended in HF for analysis. A CytoFLEX LX cytometer (Beckman Coulter) was used to collect data. Compensation was performed using the cytometer software (CytExpert v2.3) and gating was performed using FlowJo X. The antibodies used are provided in Sup. Table 6.

For sequencing analysis, the 10X genomics cell multiplexing oligo labeling for 3’ single cell RNA sequencing protocols (CG000391 rev. B) was followed. Briefly, cells were washed with HF and stained with each respective multiplexing oligo (10 conditions total). Cells were then washed 3 times with resuspension buffer, counted, and pooled in equal proportions in two separate tubes (5 samples/tube). Sequencing library preparation was prepared separately for each tube using the Chromium Single Cell 3’ Library and Gel Bead Kit v1.1, and sequenced with a target cell input number of 30,000 cells per chip. Data was processed using the CellRanger pipeline and cells with >1 hashing lipid were filtered prior to data analysis.

Following the suggested pipeline for quality control in Scanpy (v1.7.1), cells from each sequencing library were filtered to remove cells with total counts less than 1000 or greater than 40,000, percent mitochondrial genes less than 15, or number of genes by counts less than 1000 or greater than 6000. Genes expressed in less than 3 cells were removed. The two datasets were then merged and the combined dataset was normalized, and highly variable genes, PCAs, and neighbours were calculated. Cell types were named based on marker gene expression and cell counts for each cell type were calculated for each condition. These frequencies of each cell type were plotted for each Notch ligand screening condition using the matplotlib functions.

### CITE-seq and ATAC-seq data integration & analysis

Raw fastq files from all samples were aligned and quantified using CellRanger. Following the suggested pipeline for quality control in Seurat v 4.3.0 package, CITE-seq data were filtered for dead cells, doublets, and red blood cells by excluding cells with greater than 5% mitochondrial genes and less than 200, but no more than 5000 genes.

Prior to GSEA analysis, patient data (age, sex) was and then samples underwent normalization, scaling, integration using anchors, dimensional reduction, and further downstream analysis using the standard Seurat workflow with the Seurat v.4.3.0 package. PCs were visualized using an elbow plot to select dimensionality of the integrated dataset, from which we determine to implement 20 dimensions as our input for the RunUMAP and FindNeighbors clustering parameters (res = 0.9). Cells were clustered and identified based on known marker genes. To further separate cell types based on distribution of key marker genes (ex. ICOS, DPP4, MKI67), Treg cells and fibroblast cells were re-clustered by subsetting the object containing each respective cell type and then re-processing for variable features, re-scaling, and re-running FindNeighbors and FindClusters functions. Cell types were named based on expression of known marker genes and then re-added back to the main Seurat object.

Differentially expressed genes for GSEA were determined by applying model-based analysis of single-cell transcriptomics (MAST) test and the clusterProfiler package to the whole gene expression profile of male versus female for each cell type or cell type 1 versus cell type 2 for certain comparisons. The canonical pathways from the curated gene set provided by Molecular Signature Database (MsigDB) were input as the gene list for GSEA. GSEA results were outputted as a table found in supplement or visualized using the dotplot from ggplot package.

To determine common differentially expressed pathways from male and female donors, significantly (p>0.05) upregulated pathways for male and female cell types were identified. The frequency of each pathway in male and female donors was quantified to calculate the total number of cell types upregulated in each pathway and to quantify to the frequency for each main cell type classification. The top 50 most frequently upregulated pathways were plotted for male and female using the ggplot function geom_bar. Frequencies for each pathway were broken down into frequency per cell type classification.

To calculate the number of sex chromosome associated genes, we extracted a list of X and Y chromosome genes from Ensembl and calculated the percentage of differentially expressed genes (p<0.05) found on s sex chromosome for each cell type. We then calculated the mean and median number of sex chromosome-associated differentially expressed genes across cell types. The median number of genes was selected, since some cell types had few (0-3) differentially expressed genes which skewed the mean.

For integration with ATAC-seq data the general Seurat processing workflow was followed using SCTransform, which acts as a replacement for the NormalizeData, ScaleData and FindVariableFeatures functions. The data was then dimensionally reduced and filtered with RunPCA. FindNeighbors and RunUMAP were run with 30 dimensions. Cell labels were transferred from previously named cell types used for GSEA.

The ATAC-seq dat25aset was associated with the human genome (hg19) and processed using the Signac package (version 1.8.0). The dataset was filtered for peak region fragments between 750 and 80000, percent reads in peaks larger than 15, a blacklist ratio less than 0.8, a nucleosome signal less than 4 and a Transcription Start Site (TSS) Enrichment Score above 1. All filtering values were determined using Violin Plots and intercept lines. The dataset was then normalized, dimensionally reduced and clustered. To name the ATAC-seq object cell clusters, transfer anchors were identified between the ATAC-seq (query) and CITE-seq (reference) datasets. Labels were transferred to the ATAC-seq object and assigned accordingly.

CITE-seq and ATCA-seq datasets were merged and integrated using Seurat. The cell identities were reassigned according to CITE-seq labels, and the combined object was processed using the workflow described above. As the object contains multimodal data, FindMultiModalNeighbors and RunUMAP were used for final dimensional reduction and plotting. The function LinkPeaks was used to link chromatin accessibility peaks to the genes in the object.

Coverage plots were generated using the CoveragePlot function outlined in the Signac package (version 1.8.0) to visualize gene expression data in combination with chromatin accessibility. The combined ATAC-seq and CITE-seq dataset was used. The CellChat R package (version 1.6.0) was used to infer intercellular communication within the processed CITE-seq dataset. The CellChat pipeline was followed as in^46^ and the CellChat database, CellChatDB, in human was used. After processing steps, the cluster labels were re-ordered and plots were generated using built-in functions for all cell-cell interaction plots and ligand-receptor pair contribution plots. The combined ATAC-seq and CITE-seq dataset, as well as separated male and female datasets, were used in this analysis.

The pySCENIC pipeline was used to compute regulons^73^. Co-expression modules were inferred with Grnboost2, and hg19 transcription factor motif enrichment data from the Aerts laboratory was used to identify direct transcription factor targets. The area under the recovery curve metric was computed to quantify regulon activity in each cell in the dataset.

### Histology sample preparation

During thymic tissue dissociation incubations fresh-frozen CODEX tissue blocks were prepared. Tissue was rinsed in PBS, dried carefully, and placed in label histology blocks filled with O.C.T (Thermo Fisher Scientific). Blocks were frozen in cold isopentane cooled via liquid nitrogen and placed directly on dry ice once frozen. Tissue blocks were transferred to an airtight container and stored at −80°C until use.

### CODEX staining and image acquisition

CODEX buffers were described as in Schurch et al. 2020^22^ and DNA-barcoded antibodies were produced as outlined in Wang et al. 2022^23^. 8μm tissue sections were prepared, stained and imaged as described in Wang et al. 2022^23^ using an Akoya Biosciences Phenocycler microfluidics instrument.

### CODEX image analysis

Multiplex images are first processed using the custom image analysis pipeline Convolutional Registration of Images with Subpixel Precision (CRISP)^23^. Briefly, CRISP creates a multidimensional image stack by performing tile alignment and 3D-drift compensation on tile z-stack images. Images are then deconvolved and resolution-corrected through point spread function analyses, and tiles are registered and stitched. Individual cells are then detected via convolutional neural-network driven cell segmentation^23^. After cell segmentation the data can be synthesized into a matrix file resembling single cell RNA sequencing data, where individual cells are assigned quantitative expression values of all proteins captured during imaging. Using the custom image analysis pipeline HFcluster^23^, autofluorescent cells are removed and the remaining cells are clustered based on protein marker expression via Leiden based clustering. Cell types are labeled based on phenotypic expression of marker proteins as well as localization within the tissue. Finally, we perform neighbourhood analysis to detect and quantify niches within the tissue. Using K-means neighbourhood clustering, we score each cell based on the composition of its 80 nearest neighbours. Cells are then clustered into tissue neighbourhoods based on this composition score and frequency and composition of niches are quantified and compared across treatment groups.

## Supplemental Table Legends

Table 1: Postnatal thymus sample metadata.

Table 2: TotalSeq-C Human Universal Antibody Panel v1.0 used to phenotype cells via CITE-seq.

Table 3: Key marker genes defining cell types identified via CITE-seq.

Table 4: Comparison of statistically significant differentially expressed genes to sex chromosome genes.

Table 5: Cytokine concentrations used in differentiations.

Table 6: Antibodies used in Notch Ligand in vitro screening experiments.

## References

1. Spits, H. (2002). Development of αβ T cells in the human thymus. Nat. Rev. Immunol. 2, 760–772. 10.1038/nri913.

2. Lavaert, M., Valcke, B., Vandekerckhove, B., Leclercq, G., Liang, K.L., and Taghon, T. (2020). Conventional and Computational Flow Cytometry Analyses Reveal Sustained Human Intrathymic T Cell Development From Birth Until Puberty. Front. Immunol. 11.

3. Petrie, H.T., and Zúñiga-Pflücker, J.C. (2007). Zoned out: functional mapping of stromal signaling microenvironments in the thymus. Annu Rev Immunol 25, 649–679.

4. Takahama, Y. (2006). Journey through the thymus: stromal guides for T-cell development and selection. Nat. Rev. Immunol. 6, 127–135. 10.1038/nri1781.

5. Shah, D.K., and Zúñiga-Pflücker, J.C. (2014). An overview of the intrathymic intricacies of T cell development. J. Immunol. Baltim. Md 1950 *192*, 4017–4023. 10.4049/jimmunol.1302259.

6. Griffith, A.V., Fallahi, M., Nakase, H., Gosink, M., Young, B., and Petrie, H.T. (2009). Spatial Mapping of Thymic Stromal Microenvironments Reveals Unique Features Influencing T Lymphoid Differentiation. Immunity 31, 999–1009. 10.1016/j.immuni.2009.09.024.

7. Park, J.-E., Botting, R.A., Conde, C.D., Popescu, D.-M., Lavaert, M., Kunz, D.J., Goh, I., Stephenson, E., Ragazzini, R., Tuck, E., et al. (2020). A cell atlas of human thymic development defines T cell repertoire formation. Science 367, eaay3224. 10.1126/science.aay3224.

8. Bautista, J.L., Cramer, N.T., Miller, C.N., Chavez, J., Berrios, D.I., Byrnes, L.E., Germino, J., Ntranos, V., Sneddon, J.B., Burt, T.D., et al. (2021). Single-cell transcriptional profiling of human thymic stroma uncovers novel cellular heterogeneity in the thymic medulla. Nat. Commun. 12, 1096. 10.1038/s41467-021-21346-6.

9. Gustafsson, K., Isaev, S., Kooshesh, K.A., Baryawno, N., Kokkaliaris, K.D., Severe, N., Zhao, T., Scadden, E.W., Spencer, J.A., Burns, C., et al. (2022). Thymic mesenchymal niche cells drive T cell immune regeneration. 2022.10.12.511184. 10.1101/2022.10.12.511184.

10. Cordes, M., Canté-Barrett, K., van den Akker, E.B., Moretti, F.A., Kiełbasa, S.M., Vloemans, S.A., Garcia-Perez, L., Teodosio, C., van Dongen, J.J.M., Pike-Overzet, K., et al. (2022). Single-cell immune profiling reveals thymus-seeding populations, T cell commitment, and multilineage development in the human thymus. Sci. Immunol. 7, eade0182. 10.1126/sciimmunol.ade0182.

11. Carter, J.A., Strömich, L., Peacey, M., Chapin, S.R., Velten, L., Steinmetz, L.M., Brors, B., Pinto, S., and Meyer, H.V. (2022). Transcriptomic diversity in human medullary thymic epithelial cells. Nat. Commun. 13, 4296. 10.1038/s41467-022-31750-1.

12. Billiet, L., Cock, L.D., Sanchez, G.S., Mayer, R.L., Goetgeluk, G., Munter, S.D., Pille, M., Ingels, J., Jansen, H., Weening, K., et al. (2022). Single-cell profiling identifies a spectrum of human unconventional intraepithelial T lineage cells. 2022.05.24.492634. 10.1101/2022.05.24.492634.

13. Lavaert, M., Liang, K.L., Vandamme, N., Park, J.-E., Roels, J., Kowalczyk, M.S., Li, B., Ashenberg, O., Tabaka, M., Dionne, D., et al. (2020). Integrated scRNA-Seq Identifies Human Postnatal Thymus Seeding Progenitors and Regulatory Dynamics of Differentiating Immature Thymocytes. Immunity 52, 1088–1104.e6. 10.1016/j.immuni.2020.03.019.

14. Sun, V., Sharpley, M., Kaczor-Urbanowicz, K.E., Chang, P., Montel-Hagen, A., Lopez, S., Zampieri, A., Zhu, Y., de Barros, S.C., Parekh, C., et al. (2021). The Metabolic Landscape of Thymic T Cell Development In Vivo and In Vitro. Front. Immunol. 12.

15. Roels, J., Van Hulle, J., Lavaert, M., Kuchmiy, A., Strubbe, S., Putteman, T., Vandekerckhove, B., Leclercq, G., Van Nieuwerburgh, F., Boehme, L., et al. (2022). Transcriptional dynamics and epigenetic regulation of E and ID protein encoding genes during human T cell development. Front. Immunol. 13, 960918. 10.3389/fimmu.2022.960918.

16. Liang, K.L., Roels, J., Lavaert, M., Putteman, T., Boehme, L., Tilleman, L., Velghe, I., Pegoretti, V., Van de Walle, I., Sontag, S., et al. (2023). Intrathymic dendritic cell-biased precursors promote human T cell lineage specification through IRF8-driven transmembrane TNF. Nat. Immunol., 1–13. 10.1038/s41590-022-01417-6.

17. Cao, J., O’Day, D.R., Pliner, H.A., Kingsley, P.D., Deng, M., Daza, R.M., Zager, M.A., Aldinger, K.A., Blecher, R., Zhang, F., et al. (2020). A human cell atlas of fetal gene expression. Science 370, eaba7721. 10.1126/science.aba7721.

18. Gao, H., Cao, M., Deng, K., Yang, Y., Song, J., Ni, M., Xie, C., Fan, W., Ou, C., Huang, D., et al. (2022). The Lineage Differentiation and Dynamic Heterogeneity of Thymic Epithelial Cells During Thymus Organogenesis. Front. Immunol. 13, 805451. 10.3389/fimmu.2022.805451.

19. Heimli, M., Flåm, S.T., Hjorthaug, H.S., Trinh, D., Frisk, M., Dumont, K.-A., Ribarska, T., Tekpli, X., Saare, M., and Lie, B.A. (2023). Multimodal human thymic profiling reveals trajectories and cellular milieu for T agonist selection. Front. Immunol. 13.

20. Suo, C., Dann, E., Goh, I., Jardine, L., Kleshchevnikov, V., Park, J.-E., Botting, R.A., Stephenson, E., Engelbert, J., Tuong, Z.K., et al. (2022). Mapping the developing human immune system across organs. Science 376, eabo0510. 10.1126/science.abo0510.

21. Soerens, A.G., Künzli, M., Quarnstrom, C.F., Scott, M.C., Swanson, L., Locquiao, J.J., Ghoneim, H.E., Zehn, D., Youngblood, B., Vezys, V., et al. (2023). Functional T cells are capable of supernumerary cell division and longevity. Nature 614, 762–766. 10.1038/s41586-022-05626-9.

22. Schürch, C.M., Bhate, S.S., Barlow, G.L., Phillips, D.J., Noti, L., Zlobec, I., Chu, P., Black, S., Demeter, J., McIlwain, D.R., et al. (2020). Coordinated Cellular Neighborhoods Orchestrate Antitumoral Immunity at the Colorectal Cancer Invasive Front. Cell 182, 1341–1359.e19. 10.1016/j.cell.2020.07.005.

23. Wang, Y.X., Holbrook, C.A., Hamilton, J.N., Garoussian, J., Afshar, M., Su, S., Schürch, C.M., Lee, M.Y., Goltsev, Y., Kundaje, A., et al. (2022). A single cell spatial temporal atlas of skeletal muscle reveals cellular neighborhoods that orchestrate regeneration and become disrupted in aging. 2022.06.10.494732. 10.1101/2022.06.10.494732.

24. Stoeckius, M., Hafemeister, C., Stephenson, W., Houck-Loomis, B., Chattopadhyay, P.K., Swerdlow, H., Satija, R., and Smibert, P. (2017). Simultaneous epitope and transcriptome measurement in single cells. Nat. Methods 14, 865–868. 10.1038/nmeth.4380.

25. Buenrostro, J., Wu, B., Chang, H., and Greenleaf, W. (2015). ATAC-seq: A Method for Assaying Chromatin Accessibility Genome-Wide. Curr. Protoc. Mol. Biol. Ed. Frederick M Ausubel Al 109, 21.29.1–21.29.9. 10.1002/0471142727.mb2129s109.

26. Spoor, M.S., Radi, Z.A., and Dunstan, R.W. (2008). Characterization of Age and Gender-related Changes in the Spleen and Thymus from Control Cynomolgus Macaques Used in Toxicity Studies. Toxicol. Pathol. 36, 695–704. 10.1177/0192623308320279.

27. Ackman, J.B., Kovacina, B., Carter, B.W., Wu, C.C., Sharma, A., Shepard, J.-A.O., and Halpern, E.F. (2013). Sex Difference in Normal Thymic Appearance in Adults 20–30 Years of Age. Radiology 268, 245–253. 10.1148/radiol.13121104.

28. Pido-Lopez, J., Imami, N., and Aspinall, R. (2001). Both age and gender affect thymic output: more recent thymic migrants in females than males as they age. Clin. Exp. Immunol. 125, 409–413. 10.1046/j.1365-2249.2001.01640.x.

29. Hun, M.L., Wong, K., Gunawan, J.R., Alsharif, A., Quinn, K., and Chidgey, A.P. (2020). Gender Disparity Impacts on Thymus Aging and LHRH Receptor Antagonist-Induced Thymic Reconstitution Following Chemotherapeutic Damage. Front. Immunol. 11.

30. Dumont-Lagacé, M., St-Pierre, C., and Perreault, C. (2015). Sex hormones have pervasive effects on thymic epithelial cells. Sci. Rep. 5, 12895. 10.1038/srep12895.

31. Klein, S.L., and Flanagan, K.L. (2016). Sex differences in immune responses. Nat. Rev. Immunol. 16, 626–638. 10.1038/nri.2016.90.

32. Owen, D.L., La Rue, R.S., Munro, S.A., and Farrar, M.A. (2022). Tracking Regulatory T Cell Development in the Thymus Using Single-Cell RNA Sequencing/TCR Sequencing. J. Immunol. 209, 1300–1313. 10.4049/jimmunol.2200089.

33. Villani, A.-C., Satija, R., Reynolds, G., Sarkizova, S., Shekhar, K., Fletcher, J., Griesbeck, M., Butler, A., Zheng, S., Lazo, S., et al. (2017). Single-cell RNA-seq reveals new types of human blood dendritic cells, monocytes and progenitors. Science 356, eaah4573. 10.1126/science.aah4573.

34. Nitta, T., and Takayanagi, H. (2021). Non-Epithelial Thymic Stromal Cells: Unsung Heroes in Thymus Organogenesis and T Cell Development. Front. Immunol. 11.

35. Jahn, L., Kousa, A.I., Sikkema, L., Flores, A.E., Argyropoulos, K.V., Tsai, J., Lazrak, A., Nichols, K., Lee, N., Malard, F., et al. (2021). Dynamic structural cell responses in the thymus to acute injury, regeneration, and age. 2021.12.16.472014. 10.1101/2021.12.16.472014.

36. Nitta, T., Ota, A., Iguchi, T., Muro, R., and Takayanagi, H. (2021). The fibroblast: An emerging key player in thymic T cell selection. Immunol. Rev. 302, 68–85. 10.1111/imr.12985.

37. Mayne, B.T., Bianco-Miotto, T., Buckberry, S., Breen, J., Clifton, V., Shoubridge, C., and Roberts, C.T. (2016). Large Scale Gene Expression Meta-Analysis Reveals Tissue-Specific, Sex-Biased Gene Expression in Humans. Front. Genet. 7.

38. InanlooRahatloo, K., Liang, G., Vo, D., Ebert, A., Nguyen, I., and Nguyen, P.K. (2017). Sex-based differences in myocardial gene expression in recently deceased organ donors with no prior cardiovascular disease. PLOS ONE 12, e0183874. 10.1371/journal.pone.0183874.

39. Gershoni, M., and Pietrokovski, S. (2017). The landscape of sex-differential transcriptome and its consequent selection in human adults. BMC Biol. 15, 7. 10.1186/s12915-017-0352-z.

40. Lopes-Ramos, C.M., Chen, C.-Y., Kuijjer, M.L., Paulson, J.N., Sonawane, A.R., Fagny, M., Platig, J., Glass, K., Quackenbush, J., and DeMeo, D.L. (2020). Sex Differences in Gene Expression and Regulatory Networks across 29 Human Tissues. Cell Rep. 31, 107795. 10.1016/j.celrep.2020.107795.

41. McEvoy, C.M., Murphy, J.M., Zhang, L., Clotet-Freixas, S., Mathews, J.A., An, J., Karimzadeh, M., Pouyabahar, D., Su, S., Zaslaver, O., et al. (2022). Single-cell profiling of healthy human kidney reveals features of sex-based transcriptional programs and tissue-specific immunity. Nat. Commun. 13, 7634. 10.1038/s41467-022-35297-z.

42. Waldhorn, I., Turetsky, T., Steiner, D., Gil, Y., Benyamini, H., Gropp, M., and Reubinoff, B.E. (2022). Modeling sex differences in humans using isogenic induced pluripotent stem cells. Stem Cell Rep. 17, 2732–2744. 10.1016/j.stemcr.2022.10.017.

43. Oliva, M., Muñoz-Aguirre, M., Kim-Hellmuth, S., Wucher, V., Gewirtz, A.D.H., Cotter, D.J., Parsana, P., Kasela, S., Balliu, B., Viñuela, A., et al. (2020). The impact of sex on gene expression across human tissues. Science 369, eaba3066. 10.1126/science.aba3066.

44. Edgar, J.M., Michaels, Y.S., and Zandstra, P.W. (2022). Multi-objective optimization reveals time-and dose-dependent inflammatory cytokine-mediated regulation of human stem cell derived T-cell development. NPJ Regen. Med. 7, 1–13.

45. Yan, F., Mo, X., Liu, J., Ye, S., Zeng, X., and Chen, D. (2017). Thymic function in the regulation of T cells, and molecular mechanisms underlying the modulation of cytokines and stress signaling. Mol. Med. Rep. 16, 7175–7184. 10.3892/mmr.2017.7525.

46. Jin, S., Guerrero-Juarez, C.F., Zhang, L., Chang, I., Ramos, R., Kuan, C.-H., Myung, P., Plikus, M.V., and Nie, Q. (2021). Inference and analysis of cell-cell communication using CellChat. Nat. Commun. 12, 1088. 10.1038/s41467-021-21246-9.

47. Jaleco, A.C., Neves, H., Hooijberg, E., Gameiro, P., Clode, N., Haury, M., Henrique, D., and Parreira, L. (2001). Differential effects of Notch ligands Delta-1 and Jagged-1 in human lymphoid differentiation. J. Exp. Med. 194, 991–1002. 10.1084/jem.194.7.991.

48. Lehar, S.M., Dooley, J., Farr, A.G., and Bevan, M.J. (2005). Notch ligands Delta 1 and Jagged1 transmit distinct signals to T-cell precursors. Blood 105, 1440–1447. 10.1182/blood-2004-08-3257.

49. Van de Walle, I., De Smet, G., Gärtner, M., De Smedt, M., Waegemans, E., Vandekerckhove, B., Leclercq, G., Plum, J., Aster, J.C., Bernstein, I.D., et al. (2011). Jagged2 acts as a Delta-like Notch ligand during early hematopoietic cell fate decisions. Blood 117, 4449–4459. 10.1182/blood-2010-06-290049.

50. Van de Walle, I., Waegemans, E., De Medts, J., De Smet, G., De Smedt, M., Snauwaert, S., Vandekerckhove, B., Kerre, T., Leclercq, G., Plum, J., et al. (2013). Specific Notch receptor-ligand interactions control human TCR-αβ/γδ development by inducing differential Notch signal strength. J. Exp. Med. 210, 683–697. 10.1084/jem.20121798.

51. van de Laar, L., Coffer, P.J., and Woltman, A.M. (2012). Regulation of dendritic cell development by GM-CSF: molecular control and implications for immune homeostasis and therapy. Blood 119, 3383–3393. 10.1182/blood-2011-11-370130.

52. Mendes-da-Cruz, D.A., Lemos, J.P., Passos, G.A., and Savino, W. (2018). Abnormal T-Cell Development in the Thymus of Non-obese Diabetic Mice: Possible Relationship With the Pathogenesis of Type 1 Autoimmune Diabetes. Front. Endocrinol. 9.

53. Revest, J.M., Suniara, R.K., Kerr, K., Owen, J.J., and Dickson, C. (2001). Development of the thymus requires signaling through the fibroblast growth factor receptor R2-IIIb. J. Immunol. Baltim. Md 1950 167, 1954–1961. 10.4049/jimmunol.167.4.1954.

54. Bortolomai, I., Sandri, M., Draghici, E., Fontana, E., Campodoni, E., Marcovecchio, G.E., Ferrua, F., Perani, L., Spinelli, A., Canu, T., et al. (2019). Gene Modification and Three-Dimensional Scaffolds as Novel Tools to Allow the Use of Postnatal Thymic Epithelial Cells for Thymus Regeneration Approaches. Stem Cells Transl. Med. 8, 1107–1122. 10.1002/sctm.18-0218.

55. Parent, A.V., Russ, H.A., Khan, I.S., LaFlam, T.N., Metzger, T.C., Anderson, M.S., and Hebrok, M. (2013). Generation of functional thymic epithelium from human embryonic stem cells that supports host T cell development. Cell Stem Cell 13, 219–229. 10.1016/j.stem.2013.04.004.

56. Hun, M., Barsanti, M., Wong, K., Ramshaw, J., Werkmeister, J., and Chidgey, A.P. (2017). Native thymic extracellular matrix improves in vivo thymic organoid T cell output, and drives in vitro thymic epithelial cell differentiation. Biomaterials 118, 1–15. 10.1016/j.biomaterials.2016.11.054.

57. Appari, M., Werner, R., Wünsch, L., Cario, G., Demeter, J., Hiort, O., Riepe, F., Brooks, J.D., and Holterhus, P.-M. (2009). Apolipoprotein D (APOD) is a putative biomarker of androgen receptor function in androgen insensitivity syndrome. J. Mol. Med. Berl. Ger. 87, 623–632. 10.1007/s00109-009-0462-3.

58. Do Carmo, S., Séguin, D., Milne, R., and Rassart, E. (2002). Modulation of apolipoprotein D and apolipoprotein E mRNA expression by growth arrest and identification of key elements in the promoter. J. Biol. Chem. 277, 5514–5523. 10.1074/jbc.M105057200.

59. Gui, J., Mustachio, L.M., Su, D.-M., and Craig, R.W. (2012). Thymus Size and Age-related Thymic Involution: Early Programming, Sexual Dimorphism, Progenitors and Stroma. Aging Dis. 3, 280–290.

60. Heng, T.S.P., Goldberg, G.L., Gray, D.H.D., Sutherland, J.S., Chidgey, A.P., and Boyd, R.L. (2005). Effects of castration on thymocyte development in two different models of thymic involution. J. Immunol. Baltim. Md 1950 175, 2982–2993. 10.4049/jimmunol.175.5.2982.

61. Xing, Y., Wang, X., Jameson, S.C., and Hogquist, K.A. (2016). Late stages of T cell maturation in the thymus involve NF-κB and tonic type I interferon signaling. Nat. Immunol. 17, 565–573. 10.1038/ni.3419.

62. Scollay, R., Jacobs, S., Jerabek, L., Butcher, E., and Weissman, I. (1980). T cell maturation: thymocyte and thymus migrant subpopulations defined with monoclonal antibodies to MHC region antigens. J. Immunol. Baltim. Md 1950 124, 2845–2853.

63. Zelenka, T., and Spilianakis, C. SATB1-mediated chromatin landscape in T cells. Nucleus 11, 117–131. 10.1080/19491034.2020.1775037.

64. Watanabe, N., Wang, Y.-H., Lee, H.K., Ito, T., Wang, Y.-H., Cao, W., and Liu, Y.-J. (2005). Hassall’s corpuscles instruct dendritic cells to induce CD4+CD25+ regulatory T cells in human thymus. Nature 436, 1181–1185. 10.1038/nature03886.

65. Wang, J., Sekai, M., Matsui, T., Fujii, Y., Matsumoto, M., Takeuchi, O., Minato, N., and Hamazaki, Y. (2019). Hassall’s corpuscles with cellular-senescence features maintain IFNα production through neutrophils and pDC activation in the thymus. Int. Immunol. 31, 127–139. 10.1093/intimm/dxy073.

66. Tan, J., Wang, Y., Zhang, N., and Zhu, X. (2016). Induction of epithelial to mesenchymal transition (EMT) and inhibition on adipogenesis: Two different sides of the same coin? Feasible roles and mechanisms of transforming growth factor β1 (TGF-β1) in age-related thymic involution. Cell Biol. Int. 40, 842–846. 10.1002/cbin.10625.

67. Castañeda, J., Hidalgo, Y., Sauma, D., Rosemblatt, M., Bono, M.R., and Núñez, S. (2021). The Multifaceted Roles of B Cells in the Thymus: From Immune Tolerance to Autoimmunity. Front. Immunol. 12.

68. Martinez, R.J., Breed, E.R., Worota, Y., Ashby, K.M., Vobořil, M., Mathes, T., Salgado, O.C., O’Connor, C.H., Kotenko, S.V., and Hogquist, K.A. (2023). Type III interferon drives thymic B cell activation and regulatory T cell generation. Proc. Natl. Acad. Sci. 120, e2220120120. 10.1073/pnas.2220120120.

69. Lubrano di Ricco, M., Ronin, E., Collares, D., Divoux, J., Grégoire, S., Wajant, H., Gomes, T., Grinberg-Bleyer, Y., Baud, V., Marodon, G., et al. (2020). Tumor necrosis factor receptor family costimulation increases regulatory T-cell activation and function via NF-κB. Eur. J. Immunol. 50, 972–985. 10.1002/eji.201948393.

70. Owen, D.L., Mahmud, S.A., Sjaastad, L.E., Williams, J.B., Spanier, J.A., Simeonov, D.R., Ruscher, R., Huang, W., Proekt, I., Miller, C.N., et al. (2019). Thymic regulatory T cells arise via two distinct developmental programs. Nat. Immunol. 20, 195–205. 10.1038/s41590-018-0289-6.

71. Kohlhaas, S., Garden, O.A., Scudamore, C., Turner, M., Okkenhaug, K., and Vigorito, E. (2009). Cutting edge: the Foxp3 target miR-155 contributes to the development of regulatory T cells. J. Immunol. Baltim. Md 1950 *182*, 2578–2582. 10.4049/jimmunol.0803162.

72. Dong, J., Warner, L.M., Lin, L.-L., Chen, M.-C., O’Connell, R.M., and Lu, L.-F. (2021). miR-155 promotes T reg cell development by safeguarding medullary thymic epithelial cell maturation. J. Exp. Med. 218, e20192423. 10.1084/jem.20192423.

73. Aibar, S., González-Blas, C.B., Moerman, T., Huynh-Thu, V.A., Imrichova, H., Hulselmans, G., Rambow, F., Marine, J.-C., Geurts, P., Aerts, J., et al. (2017). SCENIC: single-cell regulatory network inference and clustering. Nat. Methods 14, 1083–1086. 10.1038/nmeth.4463.

74. Thiault, N., Darrigues, J., Adoue, V., Gros, M., Binet, B., Perals, C., Leobon, B., Fazilleau, N., Joffre, O.P., Robey, E.A., et al. (2015). Peripheral regulatory T lymphocytes recirculating to the thymus suppress the development of their precursors. Nat. Immunol. 16, 628–634. 10.1038/ni.3150.

75. Peligero-Cruz, C., Givony, T., Sebé-Pedrós, A., Dobeš, J., Kadouri, N., Nevo, S., Roncato, F., Alon, R., Goldfarb, Y., and Abramson, J. (2020). IL18 signaling promotes homing of mature Tregs into the thymus. eLife 9, e58213. 10.7554/eLife.58213.

76. Chen, L., Huang, H., Zheng, X., Li, Y., Chen, J., Tan, B., Liu, Y., Sun, R., Xu, B., Yang, M., et al. (2022). IL1R2 increases regulatory T cell population in the tumor microenvironment by enhancing MHC-II expression on cancer-associated fibroblasts. J. Immunother. Cancer 10, e004585. 10.1136/jitc-2022-004585.

77. Delacher, M., Simon, M., Sanderink, L., Hotz-Wagenblatt, A., Wuttke, M., Schambeck, K., Schmidleithner, L., Bittner, S., Pant, A., Ritter, U., et al. (2021). Single-cell chromatin accessibility landscape identifies tissue repair program in human regulatory T cells. Immunity 54, 702–720.e17. 10.1016/j.immuni.2021.03.007.

78. Chung, V.Y., Tan, T.Z., Tan, M., Wong, M.K., Kuay, K.T., Yang, Z., Ye, J., Muller, J., Koh, C.M., Guccione, E., et al. (2016). GRHL2-miR-200-ZEB1 maintains the epithelial status of ovarian cancer through transcriptional regulation and histone modification. Sci. Rep. 6, 19943. 10.1038/srep19943.

79. Imodoye, S.O., Adedokun, K.A., Muhammed, A.O., Bello, I.O., Muhibi, M.A., Oduola, T., and Oyenike, M.A. (2021). Understanding the Complex Milieu of Epithelial-Mesenchymal Transition in Cancer Metastasis: New Insight Into the Roles of Transcription Factors. Front. Oncol. 11.

80. Kritikou, J.S., Sánchez-Pascual, I., Muñoz-Miranda, J.P., Vashist, N., Wagner, A.K., Zhang, X.-M., Assarsson, E., Dominguez-Villar, M., Yagita, H., Garcia-Cozar, F., et al. (2022). Jagged1 overexpression on T cells induces thymic regulatory T cells leading to thymic involution (Immunology) 10.1101/2022.08.25.504005.

81. Iwabuchi, R., Ide, K., Terahara, K., Wagatsuma, R., Iwaki, R., Matsunaga, H., Tsunetsugu-Yokota, Y., Takeyama, H., and Takahashi, Y. (2021). Development of an Inflammatory CD14+ Dendritic Cell Subset in Humanized Mice. Front. Immunol. 12.

82. García-León, M.J., Fuentes, P., de la Pompa, J.L., and Toribio, M.L. (2018). Dynamic regulation of NOTCH1 activation and Notch ligand expression in human thymus development. Dev. Camb. Engl. 145, dev165597. 10.1242/dev.165597.

83. Nitta, T., Tsutsumi, M., Nitta, S., Muro, R., Suzuki, E.C., Nakano, K., Tomofuji, Y., Sawa, S., Okamura, T., Penninger, J.M., et al. (2020). Fibroblasts as a source of self-antigens for central immune tolerance. Nat. Immunol. 21, 1172–1180. 10.1038/s41590-020-0756-8.

84. Han, C.-K., Lee, W.-F., Hsu, C.-J., Huang, Y.-L., Lin, C.-Y., Tsai, C.-H., Huang, C.-C., Fong, Y.-C., Wu, M.-H., Liu, J.-F., et al. (2021). DPP4 reduces proinflammatory cytokine production in human rheumatoid arthritis synovial fibroblasts. J. Cell. Physiol. 236, 8060–8069. 10.1002/jcp.30494.

85. Zhang, K.-W., Liu, S.-Y., Jia, Y., Zou, M.-L., Teng, Y.-Y., Chen, Z.-H., Li, Y., Guo, D., Wu, J.-J., Yuan, Z.-D., et al. (2022). Insight into the role of DPP-4 in fibrotic wound healing. Biomed. Pharmacother. 151, 113143. 10.1016/j.biopha.2022.113143.

86. Tabib, T., Huang, M., Morse, N., Papazoglou, A., Behera, R., Jia, M., Bulik, M., Monier, D.E., Benos, P.V., Chen, W., et al. (2021). Myofibroblast transcriptome indicates SFRP2hi fibroblast progenitors in systemic sclerosis skin. Nat. Commun. 12, 4384. 10.1038/s41467-021-24607-6.

87. Soare, A., Györfi, H.A., Matei, A.E., Dees, C., Rauber, S., Wohlfahrt, T., Chen, C.-W., Ludolph, I., Horch, R.E., Bäuerle, T., et al. (2020). Dipeptidylpeptidase 4 as a Marker of Activated Fibroblasts and a Potential Target for the Treatment of Fibrosis in Systemic Sclerosis. Arthritis Rheumatol. Hoboken NJ 72, 137–149. 10.1002/art.41058.

88. Paton, D.N. (1904). The relationship of the thymus to the sexual organs: II. The influence of removal of the thymus on the growth of the sexual organs. J. Physiol. 32, 28–32. 10.1113/jphysiol.1904.sp001062.

89. Js, S., Gl, G., Mv, H., Ap, U., Sp, B., Ts, H., Br, B., Jl, M., Ma, M., Ap, C., et al. (2005). Activation of thymic regeneration in mice and humans following androgen blockade. J. Immunol. Baltim. Md 1950 175. 10.4049/jimmunol.175.4.2741.

90. Thomas, R., Wang, W., and Su, D.-M. (2020). Contributions of Age-Related Thymic Involution to Immunosenescence and Inflammaging. Immun. Ageing 17, 2. 10.1186/s12979-020-0173-8.

91. Chhatta, A., Mikkers, H.M.M., and Staal, F.J.T. (2021). Strategies for thymus regeneration and generating thymic organoids. J. Immunol. Regen. Med. 14, 100052. 10.1016/j.regen.2021.100052.

92. Shah, N.J., Mao, A.S., Shih, T.-Y., Kerr, M.D., Sharda, A., Raimondo, T.M., Weaver, J.C., Vrbanac, V.D., Deruaz, M., Tager, A.M., et al. (2019). An injectable bone marrow-like scaffold enhances T cell immunity after hematopoietic stem cell transplantation. Nat. Biotechnol. 37, 293–302. 10.1038/s41587-019-0017-2.

93. Ferrer, I., Garcia-Esparcia, P., Carmona, M., Carro, E., Aronica, E., Kovacs, G.G., Grison, A., and Gustincich, S. (2016). Olfactory Receptors in Non-Chemosensory Organs: The Nervous System in Health and Disease. Front. Aging Neurosci. 8.

94. Nilius, B., and Owsianik, G. (2011). The transient receptor potential family of ion channels. Genome Biol. 12, 218. 10.1186/gb-2011-12-3-218.

95. Martinez, V.G., Pankova, V., Krasny, L., Singh, T., Makris, S., White, I.J., Benjamin, A.C., Dertschnig, S., Horsnell, H.L., Kriston-Vizi, J., et al. (2019). Fibroblastic Reticular Cells Control Conduit Matrix Deposition during Lymph Node Expansion. Cell Rep. 29, 2810–2822.e5. 10.1016/j.celrep.2019.10.103.

96. Li, W., Kim, M.-G., Gourley, T.S., McCarthy, B.P., Sant’Angelo, D.B., and Chang, C.-H. (2005). An Alternate Pathway for CD4 T Cell Development: Thymocyte-Expressed MHC Class II Selects a Distinct T Cell Population. Immunity 23, 375–386. 10.1016/j.immuni.2005.09.002.

97. Choi, E.Y., Jung, K.C., Park, H.J., Chung, D.H., Song, J.S., Yang, S.D., Simpson, E., and Park, S.H. (2005). Thymocyte-Thymocyte Interaction for Efficient Positive Selection and Maturation of CD4 T Cells. Immunity 23, 387–396. 10.1016/j.immuni.2005.09.005.

98. Lee, Y.J., Jeon, Y.K., Kang, B.H., Chung, D.H., Park, C.-G., Shin, H.Y., Jung, K.C., and Park, S.H. (2009). Generation of PLZF+ CD4+ T cells via MHC class II–dependent thymocyte–thymocyte interaction is a physiological process in humans. J. Exp. Med. 207, 237–246. 10.1084/jem.20091519.

99. Ngo, S.T., Steyn, F.J., and McCombe, P.A. (2014). Gender differences in autoimmune disease. Front. Neuroendocrinol. 35, 347–369. 10.1016/j.yfrne.2014.04.004.

100. Ashpole, N.M., Logan, S., Yabluchanskiy, A., Mitschelen, M.C., Yan, H., Farley, J.A., Hodges, E.L., Ungvari, Z., Csiszar, A., Chen, S., et al. (2017). IGF-1 has sexually dimorphic, pleiotropic, and time dependent effects on healthspan, pathology, and lifespan. GeroScience 39, 129–145. 10.1007/s11357-017-9971-0.

101. Liu, Z., Mohan, S., and Yakar, S. (2016). Does the GH/IGF-1 axis contribute to skeletal sexual dimorphism? Evidence from mouse studies. Growth Horm. IGF Res. Off. J. Growth Horm. Res. Soc. Int. IGF Res. Soc. 27, 7–17. 10.1016/j.ghir.2015.12.004.

102. MacDonald, K.N., Hall, M.G., Ivison, S., Gandhi, S., Klein Geltink, R.I., Piret, J.M., and Levings, M.K. (2022). Consequences of adjusting cell density and feed frequency on serum-free expansion of thymic regulatory T cells. Cytotherapy 24, 1121–1135. 10.1016/j.jcyt.2022.06.006.

