## Supplemental Tables for "Sex biased human thymic architecture guides T cell development through spatially defined niches"

### Supplementary Tables

*Table 1: Postnatal thymus sample metadata.*

| Patient ID | Age (Months) | Sex | Thymus Weight (g) |
|---|---|---|---|
| T086 | 33 | M | 20.025 |
| T087 | 4 | F | 22.552 |
| T096 | 4 | M | - |
| T097 | 4 | F | 16.265 |
| T098 | 4 | M | 16.662 |
| T099 | 4 | F | 39 |
| T100 | 4 | M | 42 |

*Table 2: TotalSeq-C Human Universal Antibody Panel v1.0 used to phenotype cells via CITE-seq.*

| DNA_ID | Description | Clone | Barcode | Ensemble ID | Gene name |
|---|---|---|---|---|---|
| C0006 | anti-human CD86 | IT2.2 | GTCTTTGTCAGTGCA | ENSG00000114013 | CD86 |
| C0007 | anti-human CD274 (B7-H1, PD-L1) | 29E.2A3 | GTTGTCCGACAATAC | ENSG00000120217 | CD274 |
| C0020 | anti-human CD270 (HVEM, TR2) | 122 | TGATAGAAACAGACC | ENSG00000157873 | TNFRSF14 |
| C0023 | anti-human CD155 (PVR) | SKII.4 | ATCACATCGTTGCCA | ENSG00000073008 | PVR |
| C0024 | anti-human CD112 (Nectin-2) | TX31 | AACCTTCCGTCTAAG | ENSG00000130202 | NECTIN2 |
| C0026 | anti-human CD47 | CC2C6 | GCATTCTGTCACCTA | ENSG00000196776 | CD47 |
| C0029 | anti-human CD48 | BJ40 | CTACGACGTAGAAGA | ENSG00000117091 | CD48 |
| C0031 | anti-human CD40 | 5C3 | CTCAGATGGAGTATG | ENSG00000101017 | CD40 |
| C0032 | anti-human CD154 | 24-31 | GCTAGATAGATGCAA | ENSG00000102245 | CD40LG |
| C0033 | anti-human CD52 | HI186 | CTTTGTACGAGCAAA | ENSG00000169442 | CD52 |
| C0034 | anti-human CD3 | UCHT1 | CTCATTGTAACTCCT | ENSG00000167286 | CD3D |
| C0046 | anti-human CD8 | SK1 | GCGCAACTTGATGAT | ENSG00000153563 | CD8A |
| C0047 | anti-human CD56 | 5.1H11 | TCCTTTCCTGATAGG | ENSG00000149294 | NCAM1 |
| C0050 | anti-human CD19 | HIB19 | CTGGGCAATTACTCG | ENSG00000177455 | CD19 |
| C0052 | anti-human CD33 | P67.6 | TAACTCAGGGCCTAT | ENSG00000105383 | CD33 |
| C0053 | anti-human CD11c | S-HCL-3 | TACGCCTATAACTTG | ENSG00000140678 | ITGAX |
| C0058 | anti-human HLA-A,B,C | W6/32 | TATGCGAGGCTTATC | ENSG00000206503 | HLA-A |
| C0063 | anti-human CD45RA | HI100 | TCAATCCTTCCGCTT | ENSG00000081237 | PTPRC |
| C0064 | anti-human CD123 | 6H6 | CTTCACTCTGTCAGG | ENSG00000185291 | IL3RA |
| C0066 | anti-human CD7 | CD7-6B7 | TGGATTCCCGGACTT | ENSG00000173762 | CD7 |
| C0068 | anti-human CD105 | 43A3 | ATCGTCGAGAGCTAG | ENSG00000106991 | ENG |
| C0070 | anti-human/mouse CD49f | GoH3 | TTCCGAGGATGATCT | ENSG00000091409 | ITGA6 |
| C0071 | anti-human CD194 (CCR4) | L291H4 | AGCTTACCTGCACGA | ENSG00000183813 | CCR4 |
| C0072 | anti-human CD4 | RPA-T4 | TGTTCCCGCTCAACT | ENSG00000010610 | CD4 |

| | | | | | |
|---|---|---|---|---|---|
| C0073 | anti-mouse/human CD44 | IM7 | TGGCTTCAGGTCCTA | ENSG00000026508 | CD44 |
| C0081 | anti-human CD14 | M5E2 | TCTCAGACCTCCGTA | ENSG00000170458 | CD14 |
| C0083 | anti-human CD16 | 3G8 | AAGTTCACTCTTTGC | ENSG00000203747 | FCGR3A |
| C0085 | anti-human CD25 | BC96 | TTTGTCCTGTACGCC | ENSG00000134460 | IL2RA |
| C0087 | anti-human CD45RO | UCHL1 | CTCCGAATCATGTTG | ENSG00000081237 | PTPRC |
| C0088 | anti-human CD279 (PD-1) | EH12.2H7 | ACAGCGCCGTATTTA | ENSG00000188389 | PDCD1 |
| C0089 | anti-human TIGIT (VSTM3) | A15153G | TTGCTTACCGCCAGA | ENSG00000181847 | TIGIT |
| C0090 | Mouse IgG1, κ isotype Ctrl | MOPC-21 | GCCGGACGACATTAA | | |
| C0091 | Mouse IgG2a, κ isotype Ctrl | MOPC-173 | CTCCTACCTAAACTG | | |
| C0092 | Mouse IgG2b, κ isotype Ctrl | MPC-11 | ATATGTATCACGCGA | | |
| C0095 | Rat IgG2b, κ Isotype Ctrl | RTK4530 | GATTCTTGACGACCT | | |
| C0100 | anti-human CD20 | 2H7 | TTCTGGGTCCCTAGA | ENSG00000156738 | MS4A1 |
| C0101 | anti-human CD335 (NKp46) | 9E2 | ACAATTTGAACAGCG | ENSG00000189430 | NCR1 |
| C0124 | anti-human CD31 | WM59 | ACCTTTATGCCACGG | ENSG00000261371 | PECAM1 |
| C0134 | anti-human CD146 | P1H12 | CCTTGGATAACATCA | ENSG00000076706 | MCAM |
| C0136 | anti-human IgM | MHM-88 | TAGCGAGCCCGTATA | ENSG00000211899 | IGHM |
| C0138 | anti-human CD5 | UCHT2 | CATTAACGGGATGCC | ENSG00000110448 | CD5 |
| C0140 | anti-human CD183 (CXCR3) | G025H7 | GCGATGGTAGATTAT | ENSG00000186810 | CXCR3 |
| C0141 | anti-human CD195 (CCR5) | J418F1 | CCAAAGTAAGAGCCA | ENSG00000160791 | CCR5 |
| C0142 | anti-human CD32 | FUN-2 | GCTTCCGAATTACCG | ENSG00000143226 | FCGR2A |
| C0143 | anti-human CD196 (CCR6) | G034E3 | GATCCCTTTGTCACT | ENSG00000112486 | CCR6 |
| C0144 | anti-human CD185 (CXCR5) | J252D4 | AATTCAACCGTCGCC | ENSG00000160683 | CXCR5 |
| C0145 | anti-human CD103 (Integrin αE) | Ber-ACT8 | GACCTCATTGTGAAT | ENSG00000083457 | ITGAE |
| C0146 | anti-human CD69 | FN50 | GTCTCTTGGCTTAAA | ENSG00000110848 | CD69 |
| C0147 | anti-human CD62L | DREG-56 | GTCCCTGCAACTTGA | ENSG00000188404 | SELL |
| C0149 | anti-human CD161 | HP-3G10 | GTACGCAGTCCTTCT | ENSG00000111796 | KLRB1 |
| C0151 | anti-human CD152 (CTLA-4) | BNI3 | ATGGTTCACGTAATC | ENSG00000163599 | CTLA4 |
| C0152 | anti-human CD223 (LAG-3) | 11C3C65 | CATTTGTCTGCCGGT | ENSG00000089692 | LAG3 |
| C0153 | anti-human KLRG1 (MAFA) | SA231A2 | CTTATTTCCTGCCCT | ENSG00000139187 | KLRG1 |
| C0154 | anti-human CD27 | O323 | GCACTCCTGCATGTA | ENSG00000139193 | CD27 |

| | | | | | |
|---|---|---|---|---|---|
| C0155 | anti-human CD107a (LAMP-1) | H4A3 | CAGCCCACTGCAATA | ENSG00000185896 | LAMP1 |
| C0156 | anti-human CD95 (Fas) | DX2 | CCAGCTCATTAGAGC | ENSG00000026103 | FAS |
| C0158 | anti-human CD134 (OX40) | Ber-ACT35 (ACT35) | AACCCACCGTTGTTA | ENSG00000186827 | TNFRSF4 |
| C0159 | anti-human HLA-DR | L243 | AATAGCGAGCAAGTA | ENSG00000204287 | HLA-DRA |
| C0160 | anti-human CD1c | L161 | GAGCTACTTCACTCG | ENSG00000158481 | CD1C |
| C0161 | anti-human CD11b | ICRF44 | GACAAGTGATCTGCA | ENSG00000169896 | ITGAM |
| C0162 | anti-human CD64 | 10.1 | AAGTATGCCCTACGA | ENSG00000150337 | FCGR1A |
| C0163 | anti-human CD141 (Thrombomodulin) | M80 | GGATAACCGCGCTTT | ENSG00000178726 | THBD |
| C0164 | anti-human CD1d | 51.1 | TCGAGTCGCTTATCA | ENSG00000158473 | CD1D |
| C0165 | anti-human CD314 (NKG2D) | 1D11 | CGTGTTTGTTCCTCA | ENSG00000213809 | KLRK1 |
| C0167 | anti-human CD35 | E11 | ACTTCCGTCGATCTT | ENSG00000203710 | CR1 |
| C0168 | anti-human CD57 Recombinant | QA17A04 | AACTCCCTATGGAGG | ENSG00000109956 | B3GAT1 |
| C0170 | anti-human CD272 (BTLA) | MIH26 | GTTATTGGACTAAGG | ENSG00000186265 | BTLA |
| C0171 | anti-human/mouse/rat CD278 (ICOS) | C398.4A | CGCGCACCCATTAAA | ENSG00000163600 | ICOS |
| C0174 | anti-human CD58 (LFA-3) | TS2/9 | GTTCCTATGGACGAC | ENSG00000116815 | CD58 |
| C0176 | anti-human CD39 | A1 | TTACCTGGTATCCGT | ENSG00000138185 | ENTPD1 |
| C0179 | anti-human CX3CR1 | K0124E1 | AGTATCGTCTCTGGG | ENSG00000168329 | CX3CR1 |
| C0180 | anti-human CD24 | ML5 | AGATTCCTTCGTGTT | ENSG00000272398 | CD24 |
| C0181 | anti-human CD21 | Bu32 | AACCTAGTAGTTCGG | ENSG00000117322 | CR2 |
| C0185 | anti-human CD11a | TS2/4 | TATATCCTTGTGAGC | ENSG00000005844 | ITGAL |
| C0187 | anti-human CD79b (Igβ) | CB3-1 | ATTCTTCAACCGAAG | ENSG00000007312 | CD79B |
| C0189 | anti-human CD244 (2B4) | C1.7 | TCGCTTGGATGGTAG | ENSG00000122223 | CD244 |
| C0206 | anti-human CD169 (Sialoadhesin, Siglec-1) | 7-239 | TACTCAGCGTGTTTG | ENSG00000088827 | SIGLEC1 |
| C0214 | anti-human/mouse integrin β7 | FIB504 | TCCTTGGATGTACCG | ENSG00000139626 | ITGB7 |
| C0215 | anti-human CD268 (BAFF-R) | 11C1 | CGAAGTCGATCCGTA | ENSG00000159958 | TNFRSF13C |
| C0216 | anti-human CD42b | HIP1 | TCCTAGTACCGAAGT | ENSG00000203618 | GP1BB |
| C0217 | anti-human CD54 | HA58 | CTGATAGACTTGAGT | ENSG00000090339 | ICAM1 |
| C0218 | anti-human CD62P (P-Selectin) | AK4 | CCTTCCGTATCCCTT | ENSG00000174175 | SELP |
| C0219 | anti-human CD119 (IFN-γ R α chain) | GIR-208 | TGTGTATTCCCTTGT | ENSG00000027697 | IFNGR1 |
| C0224 | anti-human TCR α/β | IP26 | CGTAACGTAGAGCGA | | |
| C0236 | Rat IgG1, κ isotype Ctrl | RTK2071 | ATCAGATGCCCTCAT | | |

| | | | | | |
|---|---|---|---|---|---|
| C0238 | Rat IgG2a, κ Isotype Ctrl | RTK2758 | AAGTCAGGTTCGTTT | | |
| C0241 | Armenian Hamster IgG Isotype Ctrl | HTK888 | CCTGTCATTAAGACT | | |
| C0246 | anti-human CD122 (IL-2Rβ) | TU27 | TCATTTCCTCCGATT | ENSG00000100385 | IL2RB |
| C0247 | anti-human CD267 (TACI) | 1A1 | AGTGATGGAGCGAAC | ENSG00000240505 | TNFRSF13B |
| C0352 | anti-human FcεRIα | AER-37 (CRA-1) | CTCGTTTCCGTATCG | ENSG00000179639 | FCER1A |
| C0353 | anti-human CD41 | HIP8 | ACGTTGTGGCCTTGT | ENSG00000005961 | ITGA2B |
| C0355 | anti-human CD137 (4-1BB) | 4B4-1 | CAGTAAGTTCGGGAC | ENSG00000049249 | TNFRSF9 |
| C0358 | anti-human CD163 | GHI/61 | GCTTCTCCTTCCTTA | ENSG00000177575 | CD163 |
| C0359 | anti-human CD83 | HB15e | CCACTCATTTCCGGT | ENSG00000112149 | CD83 |
| C0363 | anti-human CD124 (IL-4Rα) | G077F6 | CCGTCCTGATAGATG | ENSG00000077238 | IL4R |
| C0364 | anti-human CD13 | WM15 | TTTCAACGCCCTTTC | ENSG00000166825 | ANPEP |
| C0367 | anti-human CD2 | TS1/8 | TACGATTTGTCAGGG | ENSG00000116824 | CD2 |
| C0368 | anti-human CD226 (DNAM-1) | 11A8 | TCTCAGTGTTTGTGG | ENSG00000150637 | CD226 |
| C0369 | anti-human CD29 | TS2/16 | GTATTCCCTCAGTCA | ENSG00000150093 | ITGB1 |
| C0370 | anti-human CD303 (BDCA-2) | 201A | GAGATGTCCGAATTT | ENSG00000198178 | CLEC4C |
| C0371 | anti-human CD49b | P1E6-C5 | GCTTTCTTCAGTATG | ENSG00000164171 | ITGA2 |
| C0373 | anti-human CD81 (TAPA-1) | 5A6 | GTATCCTTCCTTGGC | ENSG00000110651 | CD81 |
| C0384 | anti-human IgD | IA6-2 | CAGTCTCCGTAGAGT | ENSG00000211898 | IGHD |
| C0385 | anti-human CD18 | TS1/18 | TATTGGGACACTTCT | ENSG00000160255 | ITGB2 |
| C0386 | anti-human CD28 | CD28.2 | TGAGAACGACCCTAA | ENSG00000178562 | CD28 |
| C0389 | anti-human CD38 | HIT2 | TGTACCCGCTTGTGA | ENSG00000004468 | CD38 |
| C0390 | anti-human CD127 (IL-7Rα) | A019D5 | GTGTGTTGTCCTATG | ENSG00000168685 | IL7R |
| C0391 | anti-human CD45 | HI30 | TGCAATTACCCGGAT | ENSG00000081237 | PTPRC |
| C0393 | anti-human CD22 | S-HCL-1 | GGGTTGTTGTCTTTG | ENSG00000012124 | CD22 |
| C0394 | anti-human CD71 | CY1G4 | CCGTGTTCCTCATTA | ENSG00000072274 | TFRC |
| C0396 | anti-human CD26 | BA5b | GGTGGCTAGATAATG | ENSG00000197635 | DPP4 |
| C0407 | anti-human CD36 | 5-271 | TTCTTTGCCTTGCCA | ENSG00000135218 | CD36 |
| C0420 | anti-human CD158 (KIR2DL1/S1/S3/S5) | HP-MA4 | TATCAACCAACGCTT | ENSG00000125498 | KIR2DL1 |
| C0575 | anti-human CD49a | TS2/7 | ACTGATGGACTCAGA | ENSG00000213949 | ITGA1 |
| C0576 | anti-human CD49d | 9F10 | CCATTCAACTTCCGG | ENSG00000115232 | ITGA4 |
| C0577 | anti-human CD73 (Ecto-5'-nucleotidase) | AD2 | CAGTTCCTCAGTTCG | ENSG00000135318 | NT5E |
| C0581 | anti-human TCR Vα7.2 | 3C10 | TACGAGCAGTATTCA | | |
| C0582 | anti-human TCR Vδ2 | B6 | TCAGTCAGATGGTAT | | |

| | | | | | |
|---|---|---|---|---|---|
| C0591 | anti-human LOX-1 | 15C4 | ACCCTTTACCGAATA | ENSG00000173391 | OLR1 |
| C0592 | anti-human CD158b (KIR2DL2/L3, NKAT2) | DX27 | GACCCGTAGTTTGAT | ENSG00000243772 | KIR2DL3 |
| C0599 | anti-human CD158e1 (KIR3DL1, NKB1) | DX9 | GGACGCTTTCCTTGA | ENSG00000167633 | KIR3DL1 |
| C0830 | anti-human CD319 (CRACC) | 162.1 | AGTATGCCATGTCTT | ENSG00000026751 | SLAMF7 |
| C0845 | anti-human CD99 | 3B2/TA8 | ACCCGTCCCTAAGAA | ENSG00000002586 | CD99 |
| C0853 | anti-human CLEC12A | 50C1 | CATTAGAGTCTGCCA | ENSG00000172322 | CLEC12A |
| C0864 | anti-human CD352 (NTB-A) | NT-7 | AGTTTCCACTCAGGC | ENSG00000162739 | SLAMF6 |
| C0867 | anti-human CD94 | DX22 | CTTTCCGGTCCTACA | ENSG00000134539 | KLRD1 |
| C0894 | anti-human Ig light chain κ | MHK-49 | AGCTCAGCCAGTATG | ENSG00000211592 | IGKC |
| C0896 | anti-human CD85j (ILT2) | GHI/75 | CCTTGTGAGGCTATG | ENSG00000104972 | LILRB1 |
| C0897 | anti-human CD23 | EBVCS-5 | TCTGTATAACCGTCT | ENSG00000104921 | FCER2 |
| C0898 | anti-human Ig light chain λ | MHL-38 | CAGCCAGTAAGTCAC | | |
| C0902 | anti-human CD328 (Siglec-7) | 6-434 | CTTAGCATTTCACTG | ENSG00000168995 | SIGLEC7 |
| C0912 | anti-human GPR56 | CG4 | GCCTAGTTTCCGTTT | ENSG00000205336 | ADGRG1 |
| C0918 | anti-human HLA-E | 3D12 | GAGTCGAGAAATCAT | ENSG00000204592 | HLA-E |
| C0920 | anti-human CD82 | ASL-24 | TCCCACTTCCGCTTT | ENSG00000085117 | CD82 |
| C0944 | anti-human CD101 (BB27) | BB27 | CTACTTCCCTGTCAA | ENSG00000134256 | CD101 |
| C1046 | anti-human CD88 (C5aR) | S5/1 | GCCGCATGAGAAACA | ENSG00000197405 | C5AR1 |
| C1052 | anti-human CD224 | KF29 | CTGATGAGATGTCAG | ENSG00000100031 | GGT1 |

*Table 3: Key marker genes defining cell types identified via CITE-seq.*

| **Cell Type** | **Key Marker Genes** |
|---|---|
| ETPs | SELL$^{+}$, CD7$^{high}$, CD34$^{+}$ |
| Pro-T cells | TCF7$^{high}$, RAG1$^{+}$, RAG2$^{+}$, BCL11B$^{+}$, MKI67$^{low}$ |
| B-selection cells | PTCRA$^{high}$ |
| DP (P) | CD4$^{+}$, CD8$^{+}$, MKI67$^{+}$ |
| DP1 (Q) | CD4$^{+}$, CD8$^{+}$, MKI67$^{+}$, SATB1$^{high}$, CD3E$^{high}$, RAG1$^{high}$ |
| DP2 (Q) | CD4$^{+}$, CD8$^{+}$, MKI67$^{+}$, CD7$^{low}$ |
| DP-SP Transition | CD4$^{+}$, CD8$^{+}$, SATB1$^{high}$, PTCRA$^{-}$ |
| CD8 SPs | CD8$^{+}$, CD3$^{+}$ |
| CD4 SPs | CD4$^{+}$, CD3$^{+}$ |
| Pro-Treg | CD4$^{+}$CD25$^{lo}$FOXP3$^{lo}$ICOS$^{-}$CD39$^{-}$CD73$^{-}$CXCR3$^{-}$ |
| tTreg | CD4$^{+}$CD25$^{med}$FOXP3$^{med}$ICOS$^{lo}$CD39$^{-}$CD73$^{+}$CXCR3$^{-}$ |
| rrTreg | CD4$^{+}$CD25$^{hi}$FOXP3$^{hi}$ICOS$^{+}$CD39$^{+}$CD73$^{-}$CXCR3$^{+}$ |
| CD8aa T cells | CD8A$^{high}$, CD3$^{-}$ |
| Innate-like cells | NCAM1$^{+}$, CXCR3$^{low}$, CD7$^{low}$, CD56$^{low}$ |

| | |
|---|---|
| DC1 | CLEC9A$^+$, CXCR3$^+$ |
| DC1 (P) | CLEC9A$^+$, MKI67$^+$, CXCR3$^+$ |
| DC2/3 | CD1c$^+$, ITGAM$^+$, ITGAX$^+$, CD14$^+$, CD68$^+$, CD4$^+$ |
| aDCs | CCL19$^+$, LAMP3$^+$, CCR7$^+$ |
| pDCs | THBD$^+$, IL3RA$^+$, CD68$^+$, CD4$^+$, CXCR3$^+$ |
| pDCs (P) | MKI67$^+$, IL3RA$^+$, CD68$^+$, CD4$^+$, CXCR3$^+$ |
| Mono | ITGAM$^+$, ITGAX$^+$, CD14$^+$, CD68$^{low}$ |
| Mast cells | KIT$^+$ |
| B cells | CD19$^+$ |
| cTECs | PSMB11$^+$, KRT5$^+$, KRT8$^+$, EPCAM$^-$ |
| mTECs | EPCAM$^+$, KRT5$^+$, KRT8$^+$ |
| Activated mTECs | EPCAM$^+$, Class I$^{high}$, Class II$^{high}$ |
| Specialty TECs | EPCAM$^+$, NEUROD1$^+$, MYOG$^+$, POU4F1$^+$ |
| DPP4+ capFibs | DPP4$^+$, PI16$^+$, PDGFRA$^+$ |
| capFibs | DPP4$^-$, PI16$^+$, PDGFRA$^+$ |
| mFibs | PDGFRA$^+$, CCL19$^+$, Class I$^{high}$ |
| Fibs (P) | MKI67$^+$, PDGFRA$^+$ |
| KRT+ Fibs | KRT5$^+$, KRT8$^+$, PDGFRA$^+$ |
| VSMCs | MCAM$^+$, ACTA2$^+$ |
| Pericytes | MCAM$^+$, ACTA2$^-$ |
| ECs (Notch) | CD31$^+$, LYVE1$^-$, JAG1$^+$, JAG2$^+$, DLL1$^+$, DLL4$^+$ |
| ECs | CD31$^+$, LYVE1$^-$ |
| LECs | CD31$^+$, LYVE1$^+$ |

*Table 4: Comparison of statistically significant differentially expressed genes to sex chromosome genes.*

| Cell Type | Total gene count | Sex chromosome genes | Percentage of total genes |
|---|---|---|---|
| mTECs | 1976 | 69 | 3.491903 |
| cTECs | 1054 | 30 | 2.8463 |
| Activated mTECS | 430 | 18 | 4.186047 |
| Specialty TECs | 34 | 5 | 14.70588 |
| capFibs | 305 | 18 | 5.901639 |
| DPP4 capfibs | 39 | 5 | 12.82051 |
| KRT Fibs | 4 | 2 | 50 |
| mFibs | 1171 | 41 | 3.501281 |
| Fibs P | 402 | 24 | 5.970149 |
| VSMC | 81 | 6 | 7.407407 |
| Pericyte | 75 | 6 | 8 |
| ECs Notch | 24 | 3 | 12.5 |
| ECs | 166 | 13 | 7.831325 |
| LECs | 6 | 1 | 16.66667 |
| Mast | 0 | 0 | - |
| Mono | 1 | 1 | 100 |
| DC1 | 31 | 2 | 6.451613 |
| DC1 (P) | 11 | 1 | 9.090909 |

| | | | |
|---|---|---|---|
| DC2/3 | 27 | 4 | 14.81481 |
| pDCs | 6 | 0 | 0 |
| pDCs (P) | 0 | 0 | - |
| aDCs | 274 | 18 | 6.569343 |
| B cells | 315 | 12 | 3.809524 |
| Innate cells | 26 | 6 | 23.07692 |
| rrTregs | 863 | 31 | 3.592121 |
| tTregs | 511 | 17 | 3.32681 |
| Pro-Tregs | 14 | 2 | 14.28571 |
| CD8aa cells | 3 | 1 | 33.33333 |
| CD8 cells | 3743 | 117 | 3.125835 |
| CD4 cells | 179 | 8 | 4.469274 |
| DP-SP Transition | 99 | 10 | 10.10101 |
| DP1 | 924 | 40 | 4.329004 |
| DP2 | 176 | 9 | 5.113636 |
| DP (P) | 1050 | 43 | 4.095238 |
| B-selection | 6032 | 186 | 3.083554 |
| Pro-T | 523 | 30 | 5.736138 |
| ETPs | 3586 | 103 | 2.872281 |
| **Median** | | | **5.970149** |

*Table 5: Cytokine concentrations used in differentiations.*

| Cytokine | Stage 1 concentration (ng/ml) | Stage 2 concentration (ng/ml) |
|---|---|---|
| SCF | 23.9 | 120.5 |
| Flt3L | 8.7 | 8.0 |
| IL-3 | 5.3 | 1.2 |
| IL-7 | 10.0 | 44.5 |
| TNF$\alpha$ | 4.9 | 0.4 |
| CXCL12 | 9.7 | 14.8 |

*Table 6: Antibodies used in Notch Ligand in vitro screening experiments.*

| Target | Fluorochrome | Concentration | Vendor | Catalog # |
|---|---|---|---|---|
| *T-lineage* | | | | |
| CD5 | BB515 | 1:300 | eBioscience | 25-0059-42 |
| CD7 | PE-Cy7 | 1:200 | BD Biosciences | 565211 |
| *Dendritic cells* | | | | |
| CD11c | PE | 1:300 | Biolegend | 337206 |
| CD14 | VioBlue | 1:200 | Miltenyi Biotec | 130-110-582 |
| CD86 | PE-Cy7 | 1:300 | Biolegend | 374209 |
| HLA-DR | FITC | 1:300 | Biolegend | 307603 |
| *Monocyte/macrophage/mast* | | | | |

| | | | | |
|---|---|---|---|---|
| CD7 | BV605 | 1:300 | BD Biosciences | 740392 |
| CD11b | BV711 | 1:300 | Biolegend | 301343 |
| CD14 | VioBlue | 1:200 | Miltenyi Biotec | 130-110-582 |
| CD34 | APC-Cy7 | 1:200 | Biolegend | 343514 |
| CD68 | APC | 1:300 | Biolegend | 333809 |
| CD117 | BV785 | 1:200 | Biolegend | 313237 |
| FC$\varepsilon$R1$\alpha$ | PE | 1:400 | eBioscience | 12-5899-42 |
